# α/β-Hydrolase domain-containing 6 (ABHD6) accelerates the desensitization and deactivation of TARP γ-2-containing AMPA receptors

**DOI:** 10.1101/2024.06.20.599978

**Authors:** Rixu Cong, Huiran Li, Hong Yang, Jing Gu, Shanshan Wang, Qi Liu, Xiangyu Guan, Tangyunfei Su, Yulin Zheng, Dianchun Wang, Xinran Chen, Lei Yang, Yun Stone Shi, Mengping Wei, Chen Zhang

## Abstract

AMPA receptors (AMPARs) mediate most of the fast excitatory synaptic transmission in the mammalian brain. Their efficacy in responding to presynaptic glutamate release depends on their kinetics, which are determined by AMPARs and their auxiliary subunit composition. α/β-Hydrolase domain-containing 6 (ABHD6) is an AMPAR auxiliary subunit that has been shown to negatively regulate the surface delivery of AMPARs and AMPAR-mediated currents. Overexpression of ABHD6 has been shown to decrease the rising slope and increase the decay τ of mEPSCs. However, whether ABHD6 is involved in regulating AMPAR kinetics remains unclear. Here, we found that ABHD6 itself had no effect on the gating kinetics of GluA1 and GluA2(Q) containing homomeric receptors. However, in the presence of the auxiliary subunit TARP γ-2, ABHD6 accelerated the deactivation and desensitization of both GluA1 and GluA2(Q) containing homomeric receptors independent of their splicing isoforms (flip and flop) and the editing isoforms of GluA2 (R or G at position 764), except for the deactivation of GluA2(Q)i-G isoform. Besides, the recovery from desensitization of GluA1 with flip splicing isoform was slowed by the co-expression of ABHD6 in the presence of TARP γ-2. Furthermore, ABHD6 accelerated the deactivation and desensitization of GluA1i/GluA2(R)i-G and GluA2(R)i-G/GluA3(R)i heteromeric receptors in the presence of TARP γ-2. We also found that ABHD6-knockout neurons displayed slower deactivation and desensitization. Therefore, these results demonstrate that ABHD6 regulates AMPAR gating kinetics in a TARP γ-2-dependent manner.

**SIGNIFICANCE STATEMENT:** The efficacy of AMPARs in responding to presynaptic glutamate release depends on their kinetics, including deactivation, desensitization, and recovery from desensitization, which are determined by AMPARs and their auxiliary subunit composition. Using ultra-fast application of glutamate and outside-out patch recordings, we found that, in the presence of the auxiliary subunit TARP γ-2, ABHD6 accelerated the deactivation and desensitization of GluA1i/GluA2(R)i-G and GluA2(R)i-G/GluA3(R)i heteromeric receptors and GluA1 and GluA2(Q) containing homomeric receptors independent of their splicing isoforms (flip and flop) and the editing isoforms of GluA2 (R or G at position 764), except for the deactivation of GluA2(Q)i-G isoform. In ABHD6 knockout neurons, we also observed slower deactivation and desensitization. These results demonstrated that ABHD6 regulates AMPAR gating kinetics in a TARP γ-2-dependent manner.

## INTRODUCTION

The α-amino-3-hydroxy-5-methyl-4-isoxazolepropionic acid receptors (AMPARs), tetramers consisting of GluA1-4 subunits, are glutamate-gated ion channels that mediate fast excitatory synaptic transmission in the central nervous system (Bredt and Nicoll, 2003; Diering and Huganir, 2018). The transmission strength depends on the number and kinetic characteristics of AMPARs at the postsynaptic membrane (Huganir and Nicoll, 2013). AMPARs exhibit fast kinetics, with activation, deactivation, and desensitization in milliseconds, which enables the rapid depolarization of the postsynaptic membrane, ensuring a high speed and fidelity of signaling in the nervous system to accommodate the speed of information processing in the brain (Erreger et al., 2004; Mayer, 2005; Luscher and Malenka, 2012; Greger et al., 2017). AMPAR kinetics not only affects synaptic transmission and plasticity but also neurotoxicity through Ca^2+^ permeability (Huganir and Nicoll, 2013; Diering and Huganir, 2018; Yang et al., 2021). Abnormalities in these kinetics disrupt synaptic signaling and neuronal homeostasis, which are associated with a range of neurological disorders, such as Alzheimer’s disease, epilepsy, amyotrophic lateral sclerosis, and stroke (Qneibi et al., 2019; Yang et al., 2021).

In the brain, auxiliary subunits assemble with AMPARs in specific stoichiometries and modulate AMPAR kinetics (Kato et al., 2008; Brockie and Maricq, 2010; Cheng et al., 2012; Schwenk et al., 2012; Shanks NF, 2012; Yang et al., 2014; Matt et al., 2018). TARP γ-2 was the first auxiliary subunit to be discovered. It positively regulates AMPAR functions, such as promoting affinity, increasing surface expression, slowing deactivation and desensitization, and accelerating the recovery from desensitization (Gill et al., 2011a; Chen et al., 2017).

ABHD6 is a member of the α/β-hydrolase family. It was previously reported to be a serine hydrolase that could control 2-arachidonoylglycerol (2-AG) levels at cannabinoid receptors and plays a critical role in 2-AG signaling in the endocannabinoid signaling system (Chevaleyre et al., 2006; Marrs et al., 2010). High-resolution proteomic analysis has shown that ABHD6 is associated with the natural AMPAR core subunit (Schwenk et al., 2012). Functional studies revealed that overexpression of ABHD6 drastically reduces AMPAR-mediated currents by selectively inhibiting the surface expression levels of AMPARs, without affecting their total expression. The effects of ABHD6 on AMPARs were observed in both neurons and heterologous cells (Wei et al., 2016; Wei et al., 2017). Using the specific inhibitor of ABHD6 lipase activity, WWL70, or the lipase activity deficit mutation (S148A) of ABHD6, Wei et al. also found that the effects were independent of ABHD6’s hydrolase activity. Using two-dimensional gel separation of the ER and membrane fractions, Schwenk and colleagues demonstrated the assembly of AMPARs in ER and their delivery to the plasma membrane (Schwenk et al., 2019). They also showed ABHD6 trapped GluAs in the monomeric GluA-ABHD6 complex, and thus eliminated tetramer formation and surface delivery of AMPARs. In addition to this, overexpression of ABHD6 significantly increased the decay τ of mEPSCs in cultured hippocampal neurons, which indicated that ABHD6 may affect the kinetics of AMPARs (Wei et al., 2016; Wei et al., 2017). However, whether and how ABHD6 regulates AMPAR kinetics remains unknown. In this study, we investigated whether ABHD6 plays a regulatory role in the kinetic properties of functional AMPARs and whether this role is subject to AMPAR RNA editing and splice variant differences. Following previous studies (Priel et al., 2005; Milstein et al., 2007; Kato et al., 2008; Gill et al., 2011a; Khodosevich et al., 2014), we transfected different GluAs, GluAs + ABHD6, GluAs + TARP γ-2, and GluAs + TARP γ-2 + ABHD6 in HEK 293T cells and systematically studied their channel properties. Our data showed that ABHD6 did not affect the kinetic properties of AMPARs. However, in the presence of TARP γ-2, ABHD6 accelerated the deactivation and desensitization of TARP γ-2 containing GluA1 and GluA2(Q) homomeric receptors independent of their splicing isoforms and editing isoforms, except the deactivation of GluA2(Q)i-G isoform. ABHD6 also accelerated the deactivation and desensitization of GluA1i/GluA2(R)i-G and GluA2(R)i-G/GluA3(R)i heteromeric receptors. Correspondingly, ABHD6-knockout neurons exhibited slower deactivation and desensitization. Our findings revealed a new function of ABHD6 in regulating the channel properties of TARP γ-2-containing AMPA receptors.

## RESULTS

### ABHD6 reduced the glutamate-induced currents mediated by AMPARs in the presence and absence of TARP γ-2

Previous studies have shown that ABHD6 can significantly decrease currents in HEK 293T cells transfected with GluA1i, GluA2(R)i, GluA3i, GluA1i+GluA2(R)i, GluA2(R)i+GluA3i, or GluA2(R)i+GluA3i in the presence or absence of TARP γ-2 (Wei et al., 2016; Wei et al., 2017; Schwenk et al., 2019). However, whether ABHD6 inhibits other RNA editing and splice variants of AMPARs remains unclear. To answer this question, we constructed a total of 14 isoforms of GluA1, GluA2, and GluA3, specifically GluA1i, GluA1o, GluA2(Q)i-R, GluA2(Q)o-R, GluA2(Q)i-G, GluA2(Q)o-G, GluA2(R)i-R, GluA2(R)o-R, GluA2(R)i-G, GluA2(R)o-G, GluA3i-R, GluA3o-R, GluA3i-G, and GluA3o-G (illustrated in Fig. 1 A and Fig. EV1). We recorded glutamate-induced currents in HEK 293T cells transfected with these variants together with ABHD6 or control plasmid. Immunofluorescence assays and Western blot analysis were performed on cells co transfected with GluA1, TARP γ 2, and ABHD6. These experiments were conducted to verify co transfection efficiency and corresponding protein expression. Immunofluorescence results confirmed a high degree of co localization among GluA1, TARP γ 2, and ABHD6 (Fig. EV1). Our results showed that ABHD6 overexpression decreased the peak amplitudes of the currents mediated by GluA1i to ∼34%, GluA1o to ∼48%, GluA2(Q)i-R to ∼31%, GluA2(Q)o-R to ∼32%, GluA2(Q)i-G to ∼45%, and GluA2(Q)o-G to ∼21% (Fig. 1 B-C, Table. EV1.1, EV1.2). In the presence of TARP γ-2, currents were potentiated for GluA1 and GluA2 variants but not significantly for GluA3 variants, consistent with prior studies.(Pei et al., 2007; Coleman et al., 2016). Moreover, ABHD6 overexpression decreased the peak amplitudes of currents mediated by GluA1i to ∼14%, GluA1o to ∼37%, GluA2(Q)i-R to ∼9%, GluA2(Q)o-R to ∼11%, GluA2(Q)i-G to ∼6%, GluA2(Q)o-G to ∼7%, GluA2(R)i-R to ∼10%, GluA2(R)o-R to ∼44%, GluA2(R)i-G to ∼11%, GluA2(R)o-G to ∼21%, GluA3i-R to ∼5%, GluA3o-R to ∼38%, GluA3i-G to ∼21%, and GluA3o-G to ∼24% in the presence of TARP γ-2 (Fig. 1 B-D, Table. EV1.1, EV1.2). Taken together, these results showed that ABHD6 significantly reduced the glutamate-induced currents mediated by AMPARs in the presence or absence of TARP γ-2 independently of the subunit type, flip/flop splice variants, and Q/R or R/G RNA editing.

**Figure 1.**
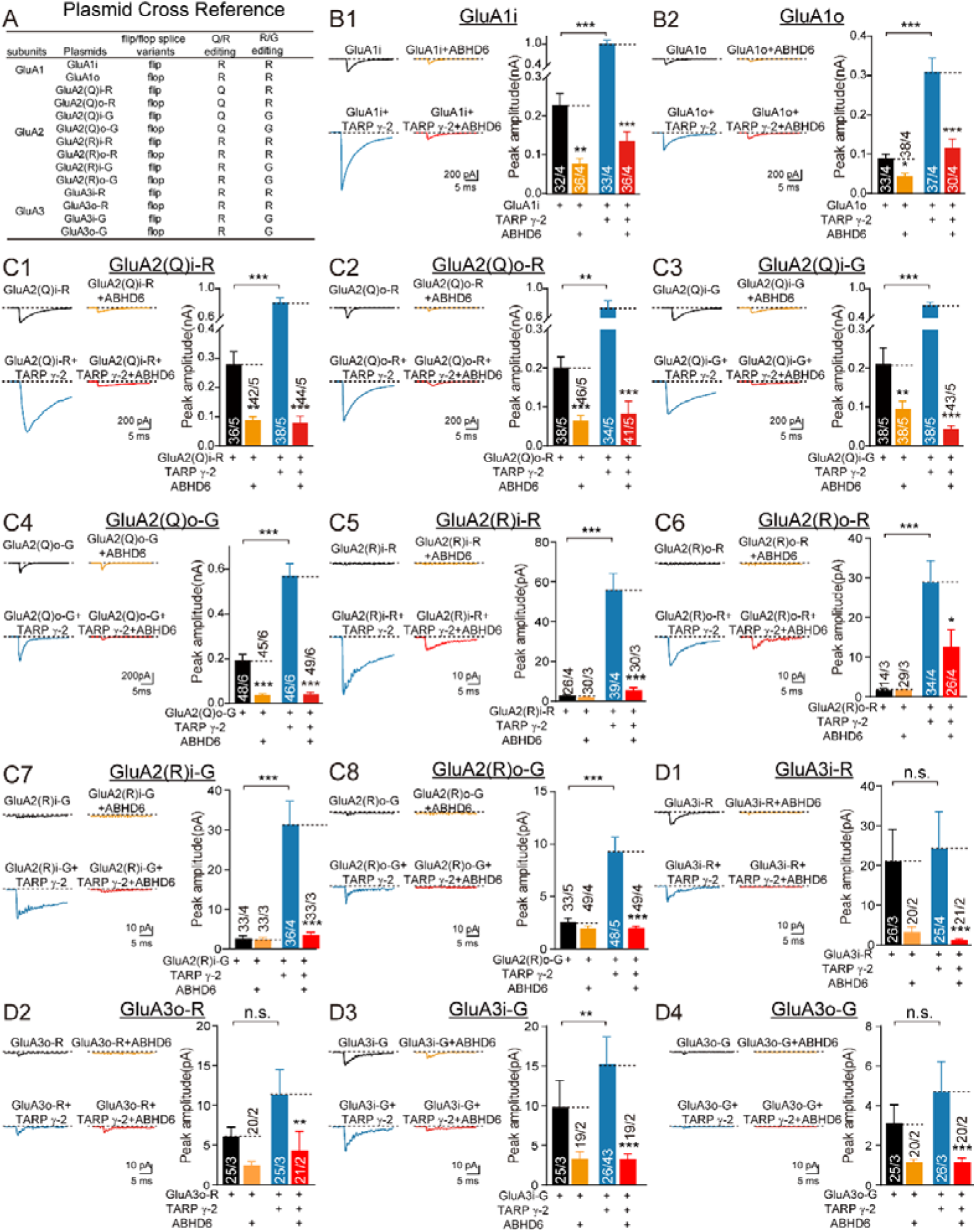
The effect of overexpression of ABHD6 on the reduction of peak current in AMPARs. (A) Plasmid abbreviations and variant combinations of AMPAR. (B-D) Representative traces (left) and summary graphs of the peak amplitudes (right) of 10 mM glutamate-induced currents in HEK 293T cells transfected with GluA1-3 (black), GluA1-3 + ABHD6 (orange), GluA1-3 + TARP γ-2 (blue), and GluA1-3 + TARP γ-2 + ABHD6 (red). The statistical method was one-way ANOVA followed by a two-way comparison (\**P* < 0.05; \*\**P* < 0.01; \*\*\**P* < 0.001. Table. EV1.2).

### ABHD6 accelerated the deactivation of homomeric AMPAR-TARP γ-2 complexes

To investigate whether ABHD6 affected AMPAR deactivation in the presence or absence of TARP γ-2, we transfected HEK 293T cells with GluAs (GluA1i, GluA1o, GluA2(Q)i-R, GluA2(Q)o-R, GluA2(Q)i-G, GluA2(Q)o-G), GluAs + ABHD6, GluAs + TARP γ-2, and GluAs + ABHD6 + TARP γ-2. To record receptor deactivation, we applied a 1-ms application of 10 mM glutamate to outside-out patches held at-60 mV. This elicited a rapidly activating inward current that decayed quickly back to baseline. Concerning the impact of GluA expression levels at the membrane on the current kinetics, we conducted Pearson correlation analyses between the amplitude and kinetics of the currents from various glutamate receptor assemblies we measured. Results showed that there is no correlation between peak amplitude and kinetics of AMPARs (Fig. EV1). Consistent with previous observations, our data showed that TARP γ-2 slowed AMPAR deactivation without selective differences in subunits, splice variants, or RNA editing (Fig. 2). Moreover, ABHD6 overexpression had a negligible effect on the deactivation of AMPARs (Fig.2 A-F, Fig. EV2 A-F, Table. EV2.1, EV2.2). However, ABHD6 overexpression reduced the deactivation time constant (τ_w, deact_) of AMPAR in the presence of TARP γ-2, except for GluA2(Q)i-G (Fig. 2 A-F, Fig. EV2 A-F, Table. EV2.1, EV2.2). Specifically, ABHD6 significantly decreased the τ_w, deact_ of GluA1i to ∼48%, of GluA1o to ∼65%, of GluA2(Q)i-R to ∼48%, of GluA2(Q)o-R to ∼60%, and of GluA2(Q)o-G to ∼60% in the presence of TARP γ-2 (Fig. 2 A-F, Fig. EV2 A-F, Table. EV2.1, EV2.2).

**Figure 2.**
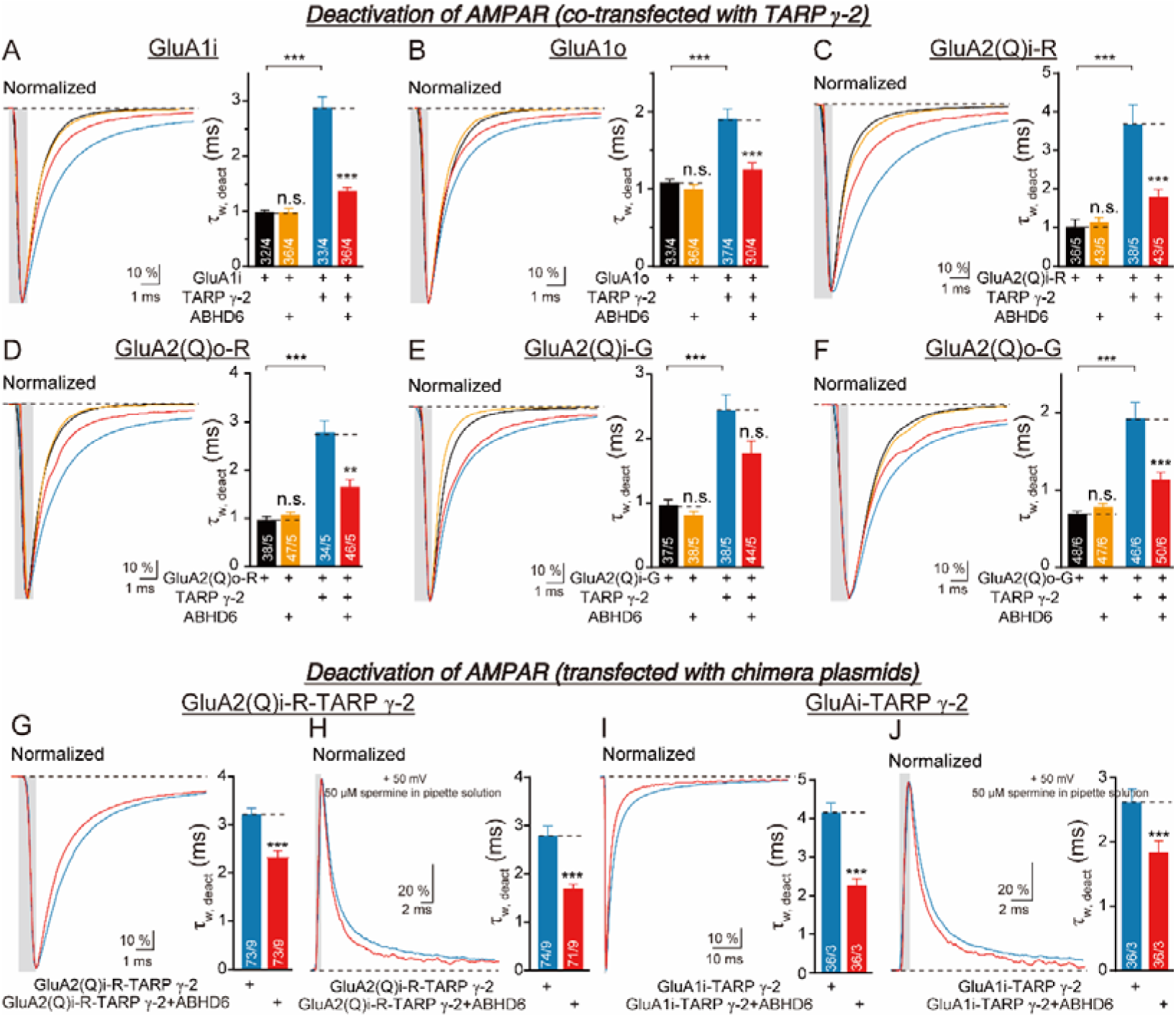
Overexpression of ABHD6 accelerated the deactivation of AMPARs-TARP. γ**-2 complexes in HEK 293T cells.** The normalized traces and the summary bar graphs of the τ _w, deact_of glutamate (10 mM Glu, 1 ms) induced currents in the outside-out patch from HEK 293T cells transfected with GluA (black), GluA + ABHD6 (orange), GluA + TARP γ-2 (blue), and GluA + TARP γ-2 + ABHD6 (red). (A) GluA1i. (B) GluA1o. (C) GluA2(Q)i-R. (D) GluA2(Q)o-R. (E) GluA2(Q)i-G. (F) GluA2(Q)o-G. (G) GluA2(Q)i-R-TARP γ-2 tandem. (I) GluA1i-TARP γ-2 tandem. γ2-containing GluA receptors could be isolated when 50 μM spermine was in the internal solution and recorded at +50 mV, the average traces and the normalized traces (right), and the summary bar graphs of the τ _w, deact_ of glutamate (10 mM Glu, 1 ms) induced currents in the outside-out patch recorded at +50 mV from HEK 293T cells transfected with GluA-TARP γ-2 tandem (blue) and GluA-TARP γ-2 tandem + ABHD6 (red). (H) GluA2(Q)i-R-TARP γ-2. (J) GluA1i-TARP γ-2. The statistical method was one-way ANOVA followed by a two-way comparison (\**P* < 0.05; \*\**P* < 0.01; \*\*\**P* < 0.001. Table. EV2.2).

We sought to determine whether the observed phenotype resulted from a direct effect of ABHD6 on TARPed AMPARs or from the changes in the ratio of TARPed to unTARPed receptors under our experimental condition. Firstly, to rule out the possibility that the observed phenotype resulted from the changes in the ratio of TARPed and unTARPed AMPARs following ABHD6 overexpression, we constructed chimeric plasmids that fused GluA2(Q)i-R or GluAi and TARP γ-2 together. We recorded the deactivation using outside-out patch recordings (-60 mV) and examined whether ABHD6 could affect its kinetics. We found that ABHD6 overexpression decreased the peak amplitudes of the currents of GluA2(Q)i-R-TARP γ-2 to ∼23% and GluA1i-TARP γ-2 to ∼25%, respectively. Concomitantly, it accelerated τ_w, deact_ mediated by GluA2(Q)i-R-TARP γ-2 to ∼65% or GluA1i-TARP γ-2 to ∼55% (Fig. 2G, 2I, Fig. EV2G, 2I, Table. EV2.1, EV2.2). These results are consistent with our previous findings using two plasmids to express AMPARs and TARP γ-2. Secondly, we isolated TARP γ-2-containing AMPA receptors using the method of Carbone et al., Nature Communications, 2016, in which they added 50 μM spermine to the internal solution and recorded at + 50 mV. We transfected chimeric plasmids GluA2(Q)i-R-TARP γ-2 or GluA1i-TARP γ-2 together with ABHD6 or control plasmid and recorded the deactivation of AMPA receptors at + 50 mV in the presence of 50 μM spermine. Our results showed that ABHD6 could still decrease the peak amplitudes of currents and accelerate the τ_w, deact_mediated by GluA2(Q)i-R-TARP γ-2 or GluA1i-TARP γ-2 (Fig. 2H, 2J, Fig. EV2H, 2J, Table. EV2.1, EV2.2). Taken together, our results clearly show that ABHD6 could affect the gating kinetics of TARPed AMPARs. Collectively, these results showed that ABHD6 accelerated the deactivation of TARP γ-2-containing AMPARs, except for the GluA2(Q)i-G variants.

### ABHD6 accelerated the desensitization of homomeric AMPAR-TARP γ-2 complexes

To investigate whether ABHD6 affected AMPAR desensitization, we recorded the desensitization induced by a 100-ms application of 10 mM glutamate using outside-out patch recordings (-60 mV). The desensitization curve showed a slower decay pattern than deactivation due to the long-term action of glutamate. Consistent with previous observations, our data showed that TARP γ-2 slowed AMPAR desensitization without selective differences in subunits, splice variants, or RNA editing (Priel et al., 2005; Coombs et al., 2012; Devi et al., 2020). ABHD6 overexpression had an insignificant effect on the desensitization of AMPARs (Fig. 3 A-F, Fig. EV3 A-F, Table. EV3.1, EV3.2). We also studied the effects of ABHD6 on the desensitization in the presence of TARP γ-2. ABHD6 overexpression reduced the desensitization time constant (τ_w, des_) of AMPARs in the presence of TARP γ-2. Specifically, it significantly decreased the τ_w, des_ of GluA1i to ∼62%, of GluA1o to ∼63%, of GluA2(Q)i-R to ∼59%, of GluA2(Q)o-R to ∼65%, of GluA2(Q)i-G to ∼60%, and of GluA2(Q)o-G to ∼71% in the presence of TARP γ-2 (Fig. 3 A-F, Fig. EV3 A-F, Table. EV3.1, EV3.2).

**Figure 3.**
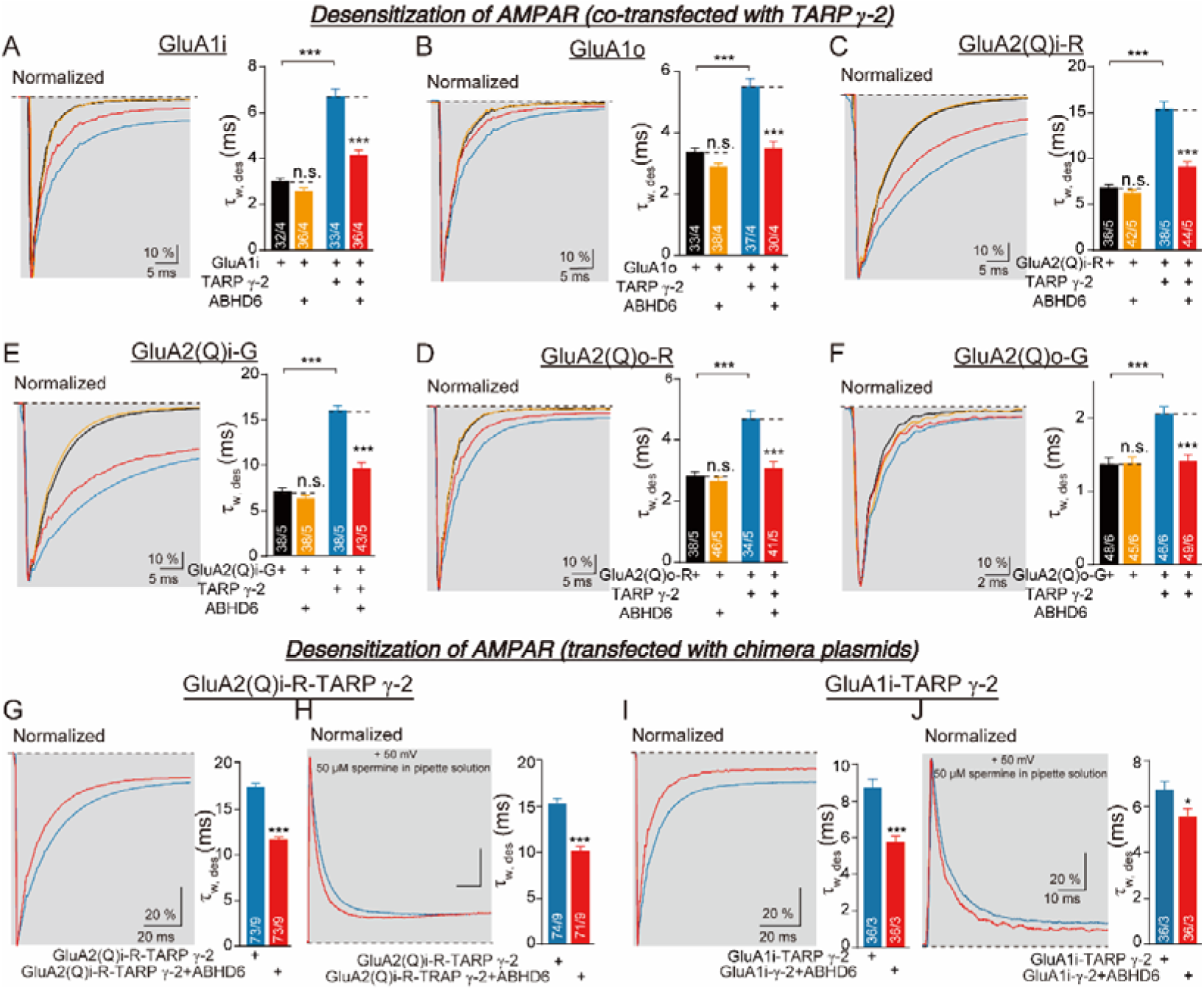
Overexpression of ABHD6 accelerated the desensitization of AMPARs-TARP. γ**-2 complexes in HEK 293T cells.** The normalized traces and the summary bar graphs of the τ _w, des_ of glutamate (10 mM Glu, 100 ms) induced currents in the outside-out patch from HEK 293T cells transfected with GluA (black), GluA + ABHD6 (orange), GluA + TARP γ-2 (blue), and GluA + TARP γ-2 + ABHD6 (red). (A) GluA1i. (B) GluA1o. (C) GluA2(Q)i-R. (D) GluA2(Q)o-R. (E) GluA2(Q)i-G. (F) GluA2(Q)o-G. (G) GluA2(Q)i-R-TARP γ-2 tandem. (I) GluA1i-TARP γ-2 tandem. TARP γ-2-containing GluA receptors could be isolated when 50 μM spermine was in the internal solution and recorded at +50 mV, the average traces and the normalized traces (right), and the summary bar graphs of the τ _w, des_, and peak amplitude of glutamate (10 mM Glu, 100 ms) induced currents in the outside-out patch recorded at +50 mV from HEK 293T cells transfected with GluA-TARP γ-2 tandem (blue) and GluA-TARP γ-2 tandem + ABHD6 (red). (H) GluA2(Q)i-R-TARP γ-2. (J) GluA1i-TARP γ-2. The statistical method was one-way ANOVA followed by a two-way comparison (\**P* < 0.05; \*\**P* < 0.01; \*\*\**P* < 0.001. Table. EV3.2).

To further confirm that the observed phenotype of ABHD6 on AMPAR desensitization was resulted from its effect on TARPed AMPARs, we also tried the chimeric plasmids that fused GluA2(Q)i-R or GluAi and TARP γ-2 together as we did in deactivation studies. We found that overexpression of ABHD6 accelerated τ_w, des_ mediated by GluA2(Q)i-R-TARP γ-2 or GluA1i-TARP γ-2 (Figure 3G, 3I, Figure EV3G, 3I, Table. EV3.1, EV3.2). Besides, we added 50 μM spermine to the internal solution and recorded at + 50 mV to isolated TARPed receptors and found that ABHD6 could still accelerate the τ_w, des_ mediated by GluA2(Q)i-R-TARP γ-2 or GluA1i-TARP γ-2 (Fig. 3H, 3J, Figure EV3H, 3J, Table. EV3.1, EV3.2).

Collectively, these results showed that ABHD6 accelerated the desensitization of TARP γ-2-containing AMPARs and that the effect did not depend on differences in GluA1/2(Q) subunits, flip/flop splice variants, or R/G editing.

### ABHD6 slowed the recovery of homomeric GluA1i-TARP γ-2 complexes from desensitization

The recovery from desensitization is another important parameter of AMPAR channel kinetics. To investigate whether ABHD6 affected AMPAR recovery from desensitization, we recorded the recovery curve in the presence or absence of TARP γ-2. Using outside-out patch recordings (-60 mV), we performed 100-ms paired-pulse stimulation at different intervals (stimulation was performed at intervals of 1-601 ms) to plot the recovery ratio curve. Consistent with previous studies, our data showed that

TARP γ-2 accelerated GluA1 recovery from desensitization (Fig. 4 A-B, Fig. EV4 A-B, Table. EV4.1, EV4.2). Moreover, it accelerated recovery of GluA2(Q)i but not GluA2(Q)o (Fig. 4 C-F, Fig. EV4 C-F, Table. EV4.1, EV4.2). We also found that ABHD6 overexpression had little effect on the recovery of AMPARs in the presence or absence of TARP γ-2, except for GluA1i-TARP γ-2, whose recovery from desensitization time constant (τ_rec_) increased to ∼157% (Fig. 4 A, Fig. EV4 A, Table. EV4.1, EV4.2). We confirmed these results using chimeric GluA2(Q)i-R-TARP γ-2 and GluA1i-TARP γ-2 plasmids, and found that ABHD6 overexpression increased the τ_rec_ of GluA1i-TARP γ-2 to ∼135%, without changing the τ_rec_ of GluA2(Q)i-R-TARP γ-2 (Fig. 4 G-H, Fig. EV4 G-H, Table. EV4.1, EV4.2). These results showed that the GluA1i-TARP γ-2 complex played a specific role in the regulation of AMPAR recovery from desensitization by ABHD6. The recovery of flip splice variants was more susceptible to regulation by other auxiliary subunits.

**Figure 4.**
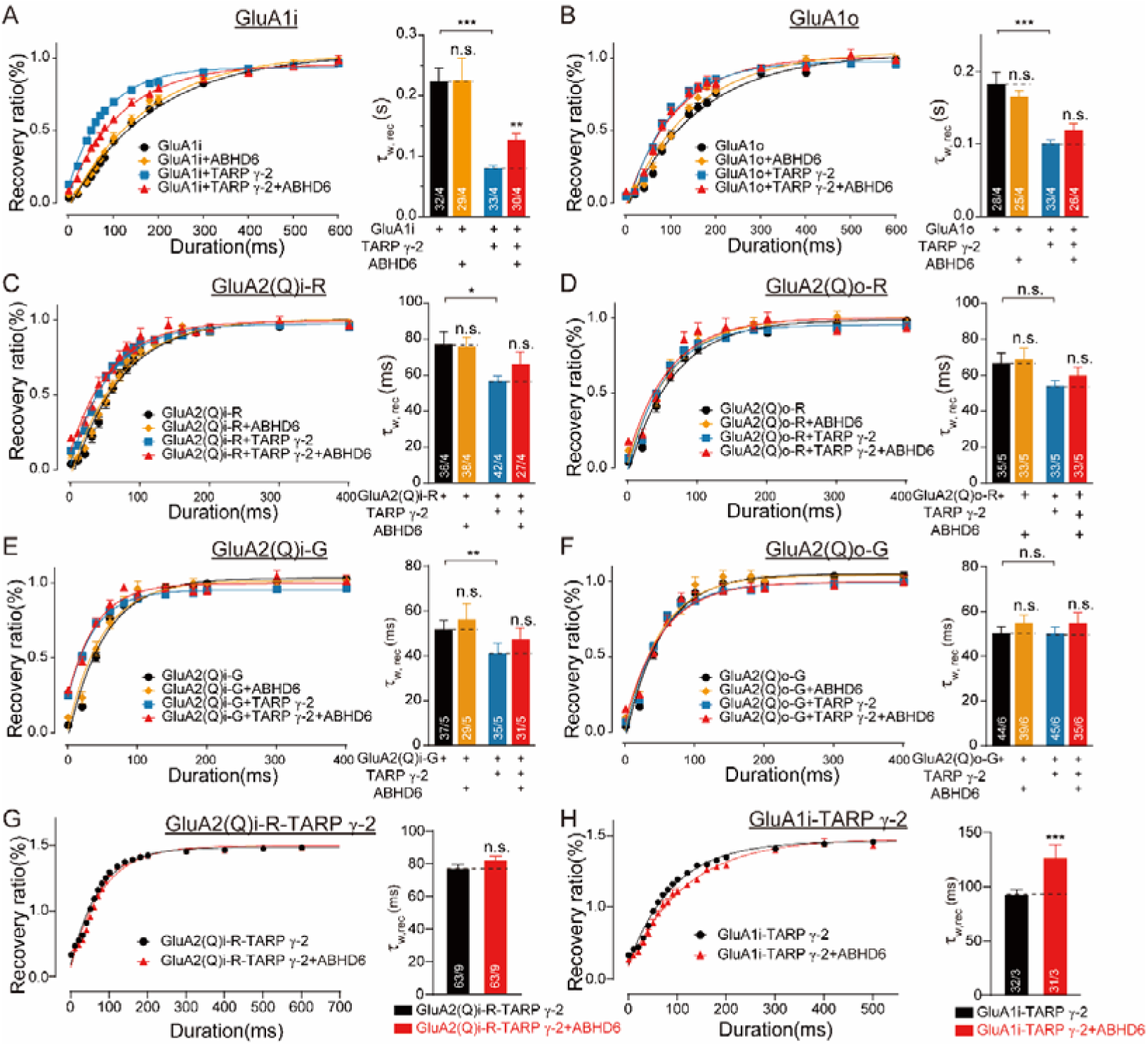
Overexpression of ABHD6 slows the recovery from desensitization of GluA1i-TARP. γ**-2 complexes in HEK 293T cells.** (A-F) Glutamate (Glu, 10 mM) induced currents in an outside-out patch excised from an HEK 293T cell transfected with GluA (black), GluA + ABHD6 (orange), GluA + TARP γ-2 (blue), and GluA+ TARP γ-2 + ABHD6 (red). The recovery ratio curves from desensitization and the summary bar graphs of the τ _w, rec._ (A) GluA1i. (B) GluA1o. (C) GluA2(Q)i-R. (D) GluA2(Q)o-R. (E) GluA2(Q)i-G. (F) GluA2(Q)o-G. (G) GluA2(Q)i-R-TARP γ-2 tandem. (H) GluA1i-TARP γ-2 tandem. The statistical method was one-way ANOVA followed by a two-way comparison (\**P* < 0.05; \*\**P* < 0.01; \*\*\**P* < 0.001. Table. EV4.2).

### ABHD6 negatively regulates the kinetics of heteromeric GluA1i/GluA2(R)i-G-TARP γ-2 and GluA2(R)i-G/GluA3(R)i-TARP γ-2 complexes

To investigate whether ABHD6 affected kinetic properties of native receptor subtypes in near-physiological states, we performed additional experiments using heteromeric receptors. Previous studies have shown that diheteromeric GluA1/GluA2 and GluA2/GluA3 receptors are the most common sets in the mammalian brain, especially in the mouse hippocampus, where the diheteromeric GluA1/GluA2 occupies more than 50% of the calcium-impermeable AMPA receptor complexes (Wenthold et al., 1996; Lu et al., 2009; Zhao et al., 2019; Yu et al., 2021). Following previous studies (Cho et al., 2007; Schwenk et al., 2009; Schwenk et al., 2012; Herring et al., 2013; Klaassen et al., 2016), we co-transfected GluA1i/GluA2(R)i-G (GluA2(R)i-G/GluA3(R)i), GluA1i/GluA2(R)i-G (GluA2(R)i-G/GluA3(R)i) + ABHD6, GluA1i/GluA2(R)i-G (GluA2(R)i-G/GluA3(R)i) + TARP γ-2, GluA1i/GluA2(R)i-G (GluA2(R)i-G/GluA3(R)i) + TARP γ-2 + ABHD6 in HEK 293T cells (GluA1i/ GluA2(R)i-G: GluA2(R)i-G/ GluA3(R)i): TARP γ-2: ABHD6 = 0.4 μg: 0.4 μg: 1.2μg: 30 ng). And we recorded the deactivation and desensitization and recovery from desensitization as previously described using outside-out patch recordings (-60 mV). Consistent with previous studies (Schwenk et al., 2009; Coombs et al., 2012), our data showed that TARP γ-2 increased the peak amplitude of currents mediated by GluA1i/GluA2(R)i-G and GluA2(R)i-G/GluA3(R)i receptors, slowed the deactivation and desensitization of GluA1i/GluA2(R)i-G and GluA2(R)i-G/GluA3(R)i receptors, but only accelerated the recovery from desensitization of GluA1i/GluA2(R)i-G receptors (Fig. 5, Fig. EV5.1, Table. EV5.1, EV5.2, EV6.1, EV6.2). Our results showed that ABHD6 accelerated the deactivation and desensitization of TARP γ-2-containing GluA1i/GluA2(R)i-G and GluA2(R)i-G/GluA3(R)i receptors, τ_w, deact_ and τ_w, des_ of GluA1i/GluA2(R)i-G–TARP γ-2 to ∼62% and ∼55%; τ_w, deact_ and τ_w, des_ of GluA2(R)i-G/GluA3(R)i–TARP γ-2 to ∼55% and ∼67%, respectively (Fig. 5A, 5B, Fig. EV5, Table. EV5.1, EV5.2). ABHD6 significantly decreased the peak amplitudes of currents mediated by GluA1i/GluA2(R)i-G receptors to ∼62%, GluA1i/GluA2(R)i-G-TARP γ-2 to ∼17%, GluA2(R)i-G/GluA3(R)i receptors to ∼32%, and GluA2(R)i-G/GluA3(R)i-TARP γ-2 to ∼13%. ABHD6 increased the τ_rec_ of GluA1i/GluA2(R)i-G and GluA1i/GluA2(R)i-G–TARP γ-2 to ∼203% and ∼198% (Fig. 5C, Fig. EV5, Table. EV6.1, EV6.2). These results showed that in the presence of TARP γ-2, ABHD6 specifically accelerates the inactivation and desensitization processes of heterodimeric AMPARs (GluA1/2 and GluA2/3). Concurrently, ABHD6 broadly reduces the current amplitude of all receptor subtypes and delays the recovery from desensitization in GluA1/2 receptors. Our results indicated that ABHD6 plays a crucial role in TARP γ-2-dependent regulation of the kinetic properties of AMPA receptors.

**Figure 5.**
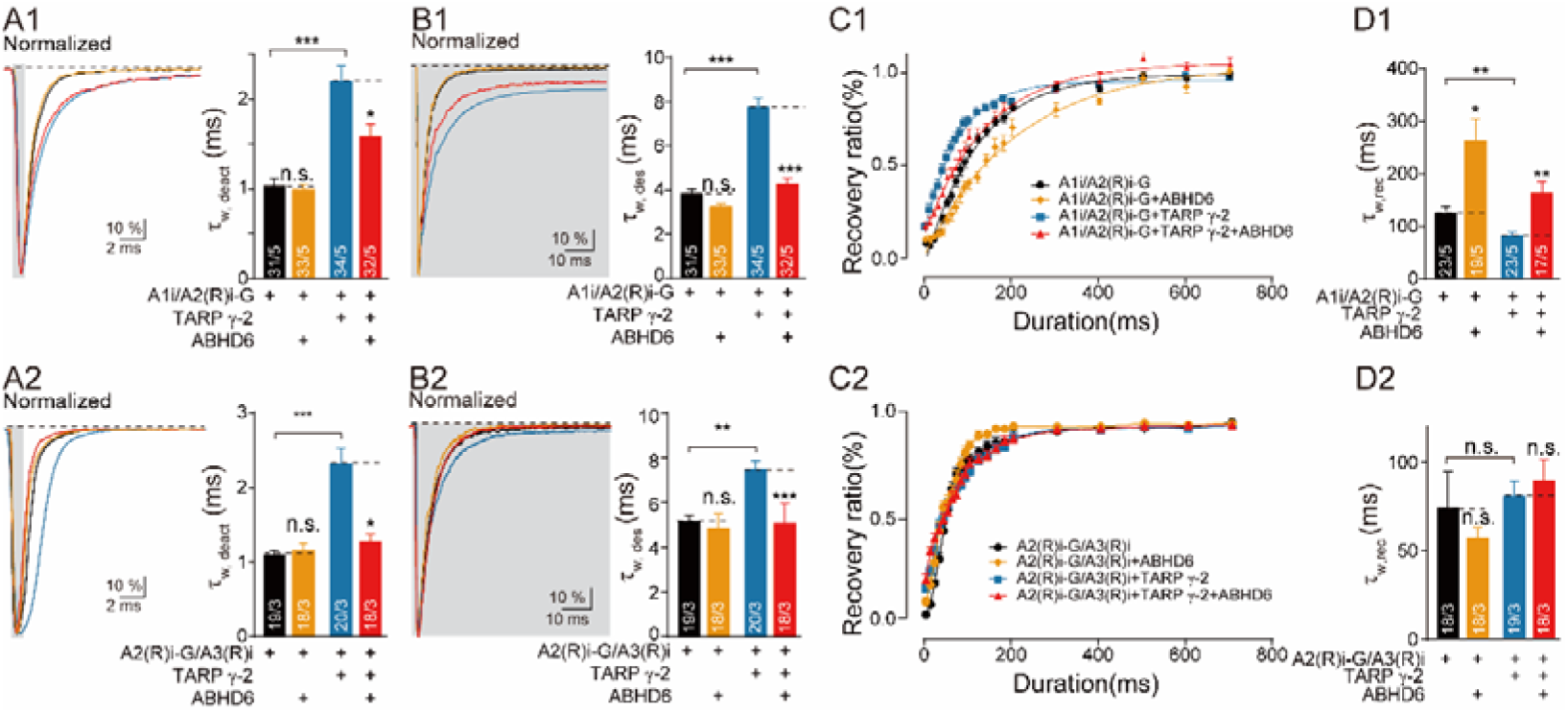
Overexpression of ABHD6 accelerated the deactivation and desensitization of GluA1i/GluA2(R)i-G (or GluA2(R)i-G/GluA3(R)i) receptors-TARP. γ**-2 complexes in HEK 293T cells and slowed the recovery of GluA1i/GluA2(R)i-G receptors in the presence and absence of TARP** γ**-2.** (A) The normalized traces and the summary bar graphs of the τ _w, deact_ of glutamate (10 mM Glu, 1 ms) induced currents in the outside-out patch recorded at-60 mV from HEK 293T cells transfected with GluA1i/GluA2(R)i-G or GluA2(R)i-G/GluA3(R)i (black), GluA1i/GluA2(R)i-G or GluA2(R)i-G/GluA3(R)i + ABHD6 (orange), GluA1i/GluA2(R)i-G or GluA2(R)i-G/GluA3(R)i + TARP γ-2 (blue), and GluA1i/GluA2(R)i-G or GluA2(R)i-G/GluA3(R)i + TARP γ-2 + ABHD6 (red). (B) The normalized traces and the summary bar graphs of the τ _w, des_, and peak amplitude of glutamate (10 mM Glu, 100 ms) induced currents in the outside-out patch recorded at-60 mV from HEK 293T cells transfected with GluA1i/GluA2(R)i-G or GluA2(R)i-G/GluA3(R)i (black), GluA1i/GluA2(R)i-G or GluA2(R)i-G/GluA3(R)i + ABHD6 (orange), GluA1i/GluA2(R)i-G or GluA2(R)i-G/GluA3(R)i + TARP γ-2 (blue), and GluA1i/GluA2(R)i-G or GluA2(R)i-G/GluA3(R)i + TARP γ-2 + ABHD6 (red). (C-D) Glutamate (Glu, 10 mM) induced currents in an outside-out patch excised from an HEK 293T cell transfected with GluA1i/GluA2(R)i-G or GluA2(R)i-G/GluA3(R)i (black), GluA1i/GluA2(R)i-G or GluA2(R)i-G/GluA3(R)i + ABHD6 (orange), GluA1i/GluA2(R)i-G or GluA2(R)i-G/GluA3(R)i + TARP γ-2 (blue), and GluA1i/GluA2(R)i-G or GluA2(R)i-G/GluA3(R)i + TARP γ-2 + ABHD6 (red). The first application of 100 ms glutamate was followed by a second glutamate application at increasing pulse intervals at-60 mV. The recovery ratio curves from desensitization _(C)_ and the summary bar graphs of the τ _w, rec_ (D) The statistical method was one-way ANOVA followed by a two-way comparison (*P < 0.05; **P < 0.01; ***P < 0.001. Table. EV5.2 and Table. EV6.2)

### ABHD6 negatively regulates the kinetics of AMPA receptors in neurons

The kinetic properties of AMPA receptors play an important regulatory role in neuronal excitability, synaptic plasticity, and neural circuit function. Having established ABHD6’s TARP γ-2-dependent function in recombinant receptors, we next asked whether this mechanism operates in neurons, where AMPARs are primarily heteromeric (GluA1/2 and GluA2/3) and embedded in diverse auxiliary protein networks (Schwenk et al., 2012). To investigate whether ABHD6 is involved in the kinetic regulation of neurons, we used ABHD6 knockout cells (brief description of the experimental process). Consistent with our model, ABHD6 knockout neurons exhibited significantly slower deactivation and desensitization kinetics, confirming the physiological relevance of this regulatory pathway. Our results showed that ABHD6 knockout slowed the deactivation and desensitization of AMPA receptors, τ_w, deact_ and τ_w, des_ to ∼121% and ∼148%. ABHD6 knockout also increased the peak amplitudes of currents mediated by AMPA receptors to ∼62% (Fig. 6, Table. EV7.1, EV7.2). These findings establish ABHD6 as a novel regulator of TARP γ-2-dependent AMPAR gating dynamics.

**Figure 6.**
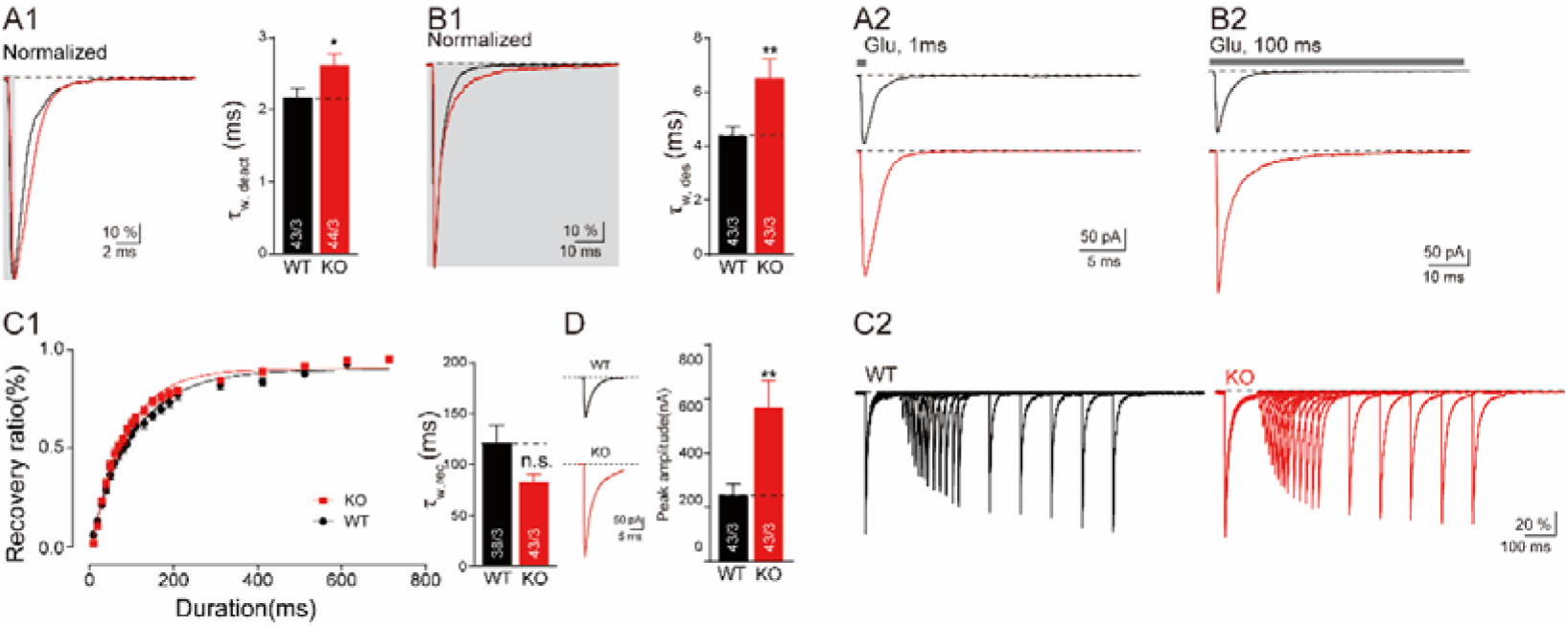
Deletion of ABHD6 decelerated the deactivation and desensitization in ABHD6 KO primary hippocampus neurons. (A) The normalized traces and the summary bar graphs of the τ _w, deact_ of Glutamate (10 mM Glu, 1 ms) induced currents in the outside-out patch recorded at-70 mV from wild-type primary hippocampus neurons (black) and ABHD6 knockout (ABHD6 KO) primary hippocampus neurons (red); (A2) the average traces of the τ _w, deact_ of Glutamate (10 mM Glu, 1 ms) induced currents in the outside-out patch recorded at - 70 mV from primary hippocampus neurons. (B) The normalized traces and the summary bar graphs of the τ _w, des_ of Glutamate (10 mM Glu, 100 ms) induced currents in the outside-out patch recorded at-70 mV from wild-type primary hippocampus neurons (black) and ABHD6 knockout (ABHD6 KO) primary hippocampus neurons (red); (B2) the average traces of the τ _w, des_ of Glutamate (10 mM Glu, 1 ms) induced currents in the outside-out patch recorded at - 70 mV from primary hippocampus neurons. (C) Glutamate (Glu, 10 mM) induced currents in an outside-out patch excised from wild-type primary hippocampus neurons (black) and ABHD6 knockout (ABHD6 KO) primary hippocampus neurons (red). The first application of 100 ms glutamate was followed by a second glutamate application at increasing pulse intervals at-70 mV. The recovery ratio curves from desensitization (C1, left), the summary bar graphs of the τ _w, rec_ (C2, right), and the typical trace of recovery from desensitization (C2). (D) The typical trace and the summary bar graphs of glutamate (10 mM Glu, 100 ms) induced currents in the outside-out patch from WT primary hippocampus neurons (black) and ABHD6 knockout (ABHD6 KO) primary hippocampus neurons (red). The statistical method was one-way ANOVA followed by a two-way comparison (*P < 0.05; **P < 0.01; ***P < 0.001. Table. EV7.2)

## DISCUSSION

In this study, we systematically study the effects of ABHD6 on the amplitudes and kinetics of AMPARs with different subunit types, including GluA1(flip/flop), GluA2(Q) (flip/flop, R/G) and diheteromeric GluA1i/GluA2(R)i-G and GluA2(R)i-G/GluA3(R)i receptors, in the presence or absence of TARP γ-2 in HEK 293T cells. Additionally, we obtained identical results in GluA4, which is expressed at higher levels in the cerebellum and brainstem (Fig.7, EV7, Table. EV8.1, EV8.2). Our results showed that ABHD6 inhibited glutamate-induced currents in all different AMPARs and regulated the gating kinetics of AMPARs in the presence of TARP γ-2. Furthermore, we validated these effects in neurons with ABHD6 knockout. These findings highlight ABHD6 as a critical regulator of AMPAR function, influencing both receptor trafficking and synaptic efficacy, with broad implications for synaptic plasticity and neuronal excitability (Wang et al., 2025).

**Figure 7.**
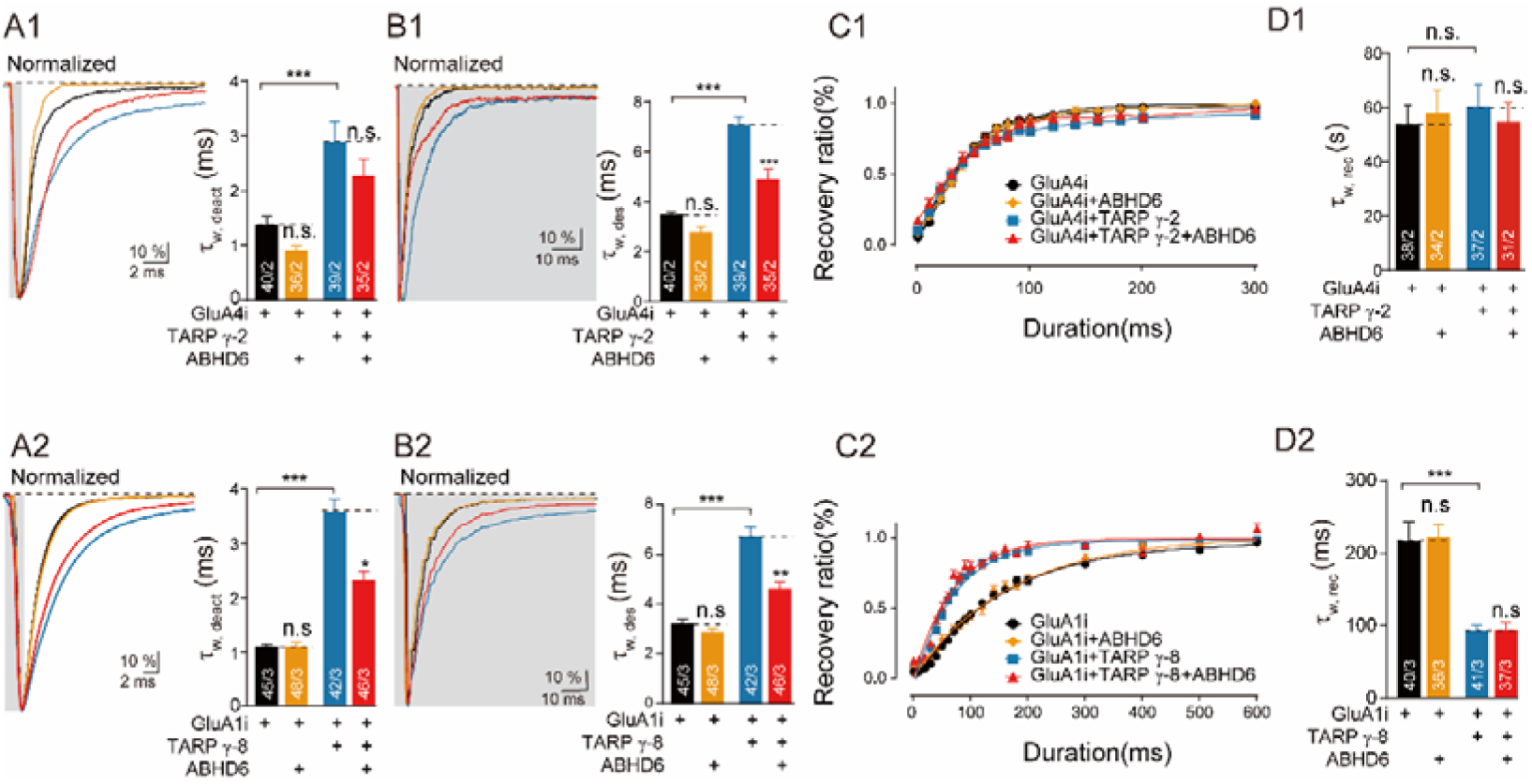
ABHD6 accelerated the deactivation and desensitization of homomeric GluA4i-TARP. γ**-2 complexes and negatively regulates the kinetics of GluA1i-TARP** γ**-8 complexes** (A) The normalized traces and the summary bar graphs of the τ _w, deact_ of glutamate (10 mM Glu, 1 ms) induced currents in the outside-out patch recorded at-60 mV from HEK 293T cells transfected with GluA4i or GluA1i (black), GluA4i or GluA1i (black) + ABHD6 (orange), GluA4i or GluA1i + TARP γ-2 (blue), or GluA4i or GluA1i + TARP γ-2 + ABHD6 (red). (B) The normalized traces and the summary bar graphs of the τ _w, des_of glutamate (10 mM Glu, 100 ms) induced currents in the outside-out patch recorded at-60 mV from HEK 293T cells transfected with GluA4i or GluA1i (black), GluA4i or GluA1i (black) + ABHD6 (orange), GluA4i or GluA1i + TARP γ-2 (blue), or GluA4i or GluA1i + TARP γ-2 + ABHD6 (red). (C-D) Glutamate (Glu, 10 mM) induced currents in an outside-out patch excised from an HEK 293T cell transfected with GluA4i or GluA1i (black), GluA4i or GluA1i (black) + ABHD6 (orange), GluA4i or GluA1i + TARP γ-2 (blue), or GluA4i or GluA1i + TARP γ-2 + ABHD6 (red). The statistical method was one-way ANOVA followed by a two-way comparison (*P < 0.05; **P < 0.01; ***P < 0.001. Table. EV8.2, EV 9.2)

Previous studies have reported that ABHD6 can reduce glutamate-induced currents and the surface expression of GluAs independently of the subunit composition (Wei et al., 2016; Wei et al., 2017). They found that ABHD6 overexpression can reduce AMPAR currents and surface expression levels of AMPARs in HEK 293T cells transfected with GluA1i, GluA2(R)i, GluA3i, GluA1i + GluA2(R)i, GluA2(R)i + GluA3i, or GluA2(R)i + GluA3i in the presence or absence of TARP γ-2. In addition to flip splice of AMPAR subunits, mammalian brains also have the flop splicing isoform. The flop splice variants are expressed in a pattern that differs from but partially overlaps with flip splice variants. The CA1 and CA3 regions of the hippocampus predominantly express flip splice variants, whereas DG highly expresses flop splice variants (Sommer et al., 1990). Flip splice variants remain unchanged in the developing brain since birth, whereas flop splice variants are expressed at low levels in the first eight days of life and reach adult levels by day 14, suggesting that flop splice variants may be involved in the formation of mature receptors (Monyer et al., 1991). Beyond subunit composition and splicing variants, the function of AMPARs is also finely regulated by RNA editing. Q/R editing enables the conversion of neutral to positively charged residues in the ion-selective filter of the channel, causing impermeability to divalent cations such as Ca2+. This not only alters channel conductance and current but also contributes to neuronal dysfunction and excitotoxicity (Kawahara et al., 2004; Kwak and Kawahara, 2004). R/G editing markedly influences receptor desensitization and recovery kinetics, and may modulate interactions with auxiliary proteins, thereby playing a critical role in synaptic plasticity and development (Stern-Bach et al., 1998; Coombs et al., 2012; Wright and Vissel, 2012; Li et al., 2025). The conversion from R to G weakens inter-dimer interactions within the binding domains, leading to structurally more flexible receptors (Lomeli et al., 1994). Furthermore, R/G editing exhibits strong developmental regulation and varies across brain regions and cell types (Geiger et al., 1995). Therefore, in this study, we systematically examined the effect of ABHD6 on different flip/flop splice variants and R/G editing subtypes. Our results demonstrate that ABHD6 also suppresses currents in HEK 293T cells expressing flop splice variants and R/G-edited receptors. Thus, this study provided further evidence that ABHD6 inhibited AMPARs on the amplitude of glutamate-induced currents. The reason for the decreased AMPAR-mediated currents might be due to the negative effects of ABHD6 on the tetramer formation of AMPARs reported by Schwenk J and colleagues (Schwenk et al., 2019).

The gating dynamics of AMPA receptors (AMPARs) are significantly regulated by multiple auxiliary subunits to finely modulate synaptic transmission(Khodosevich et al., 2014). Classic transmembrane AMPA receptor-associated proteins (TARPs γ-2, γ-3, γ-4, γ-8) generally slow the deactivation and desensitization processes (Cho et al., 2007; Milstein et al., 2007), whereas TARP γ-5 accelerates these processes (Kato et al., 2008). Other key modulators include cornichon family AMPA receptor auxiliary proteins (CNIH-2/3) and GSG1L, which generally slow receptor kinetics in heterologous expression systems (Kato et al., 2010; Schwenk et al., 2012), although their effects in neurons can be context-dependent (Gu et al., 2016; Mao et al., 2017). Additional diversity arises from synapse enriched proteins such as SynDIG4 and CKAMP44, which exert complex and sometimes opposing effects on different kinetic parameters (Matt et al., 2018; Khodosevich et al., 2014). This diversity comes from the known co-assembly of AMPA receptor subunits (the pore-forming GluA subunit) with three classes of auxiliary proteins—collectively comprising 21 components, most of which are secretory or transmembrane proteins. Importantly, multiple auxiliary subunits (e.g., TARP γ-8 and CNIH-2) can co-assemble within a single AMPAR complex, and their combined presence modulates functional outcomes in ways not predicted by individual subunits alone, underscoring a combinatorial regulatory logic (Shi et al., 2010; Yu et al., 2021; Herring et al., 2013). Given that native synaptic AMPARs predominantly exist as GluA2-containing hetero-oligomers (e.g., GluA1/2, GluA2/3), although homo-oligomers have also structurally validated, understanding how novel auxiliary proteins such as ABHD6 integrate into this complex framework becomes paramount (Lu et al., 2009; Wenthold et al., 1996; Zhao et al., 2016; Malinow and Malenka, 2002).

Against this backdrop, we systematically investigated the effect of ABHD6 on the kinetic characteristics of either homomeric AMPARs or heteromeric AMPARs. Our results clearly demonstrated that, unlike previously identified auxiliary subunits, ABHD6 accelerates GluA1 and GluA2(Q) deactivation and desensitization in the presence but not in the absence of TARP γ-2, which is also consistent with the previous publication that overexpression of ABHD6 accelerates AMPARs-mediated deactivation in neurons (Wei et al., 2017). The results may suggest a novel mechanism by which auxiliary subunits regulate the kinetics of AMPARs and their ability not to regulate the kinetics of GluAs but to alter the kinetics of native AMPARs by altering the stoichiometry of TARPs in the AMPARs complex. Moreover, its effects do not depend on the flip/flop isoform or R/G editing of AMPARs. We did not investigate the effects of ABHD6 on the kinetics of GluA2 Q607R–edited and GluA3-related AMPARs because their glutamate-induced currents are not detectable.

Although there is no direct evidence indicating that ABHD6 and TARP γ-2 bind to each other, both are known to associate with AMPA receptors, suggesting the possibility of indirect or regulatory interactions. For example, their relationship could be transient, condition-dependent, or mediated through mechanisms such as conformational changes or steric hindrance (Gill et al., 2011b; Sumioka, 2013; Wei et al., 2017). Studies have reported that scaffold proteins participate in the binding, anchoring, maintenance, and removal of AMPA receptors, either through direct interaction with receptors or through indirect binding via auxiliary subunits (Danielson et al., 2014; Chen et al., 2025). Additionally, we extended the same experimental approach to AMPA receptors containing the GluA1 flip subtype together with TARP γ-8. Our results demonstrate that this ABHD6-dependent regulatory mechanism also applies to other TARP family members, including TARP γ-8 (Figure 7, EV7, Table. EV9.1, EV9.2). Our findings indicate that ABHD6 plays a critical negative regulatory role on AMPA receptor function. It suppresses synaptic current amplitude and accelerates the deactivation and desensitization kinetics in a TARP γ-2-dependent manner. By shortening synaptic response duration and reducing total charge transfer, ABHD6 may thereby restrain neuronal excitability and narrow the temporal window for synaptic integration. Loss of ABHD6 function—as observed in our knockout neurons, which exhibit slowed kinetics—could promote excitatory hyperactivity. Thus, as a key “molecular brake” on synaptic excitability, dysregulation of ABHD6 may directly contribute to the pathogenesis of neurological disorders (Zhang et al., 2025). Insufficient braking function may lead to excessive synaptic transmission, strongly correlating with hyperexcitability conditions such as epilepsy (Liu et al., 2025). Conversely, overly potent braking might result in synaptic dysfunction, potentially contributing to early synaptic impairment in cognitive disorders like Alzheimer’s disease. Overall, our research highlights ABHD6 as a promising target for novel therapeutic strategies in neurological disorders and provides a solid theoretical foundation for further investigation in this field.

There are still questions that need to be answered. What are the molecular mechanisms behind the role of ABHD6 in regulating AMPAR kinetics in a TARP γ-2-dependent manner? The structure of the AMPARs-TARP γ-2 complex has revealed that the TMD of TARP γ-2 is arranged around the TMD of AMPARs and serves to support the ion channel pore of the receptor (Chen et al., 2017; Zhao et al., 2019). When AMPARs are activated, the differentially charged extracellular region of TARP γ-2 binds to the LBD of AMPARs, stabilizes the intra-and inter-dimeric interfaces of AMPARs, and regulates the activation, deactivation, and desensitization of AMPARs (Tomita et al., 2005; Zhao et al., 2016). Previous studies have shown the binding of ABHD6 to the CTD of AMPARs through pull-down experiments (Wei et al., 2017). However, the CTD structure of TARP γ-2 and AMPARs has not been reported. Further structure-based studies may provide more information about the channel gating properties of AMPARs.

## MATERIALS AND METHODS

### Construction of expression vectors

Cloning of mouse ABHD6-2A-GFP containing pFUGW was performed as previously described. GluA1-3 cDNA (rat) was applied to plasmid construction as previously described (Wei et al., 2017; Jiang et al., 2021). The flip and flop isoforms were cloned into an IRES-GFP expression vector using polymerase chain reaction (PCR). The flip and flop isoforms were cloned into an IRES-GFP expression vector using polymerase chain reaction (PCR). Q/R and R/G editing variants were generated by PCR-based cloning and FastCloning. GluA1 and TARP γ-2 were subcloned using EcoRI and SalI sites (Milstein et al., 2007), GluA2 and GluA3 were inserted with XhoI and SalI, and GluA4 was inserted with EcoRI and BamHI. All constructs were verified by restriction mapping and sequencing of PCR-amplified regions. In GluAs-γ-2 tandems, a linker was added between GluAs and TARP γ-2. Seventeen plasmids of GluA were constructed: GluA1i, GluA1o, GluA2(Q)i-R, GluA2(Q)o-R, GluA2(Q)i-G, GluA2(Q)o-G, GluA2(R)i-R, GluA2(R)o-R, GluA2(R)i-G, GluA2(R)o-G, GluA3i-R, GluA3o-R, GluA3i-G, GluA3o-G, GluA4i, GluA1i-γ-2, and GluA2(Q)i-R-γ-2.

### HEK 293T cell culture and transfection

HEK 293T cells were cultured in Dulbecco’s Modified Eagle Medium (Gibco) containing 10% fetal bovine serum (ExCell Bio) and 1% penicillin–streptomycin (Gibco) in an incubator with 5% CO_2_ at 37 [. The transfection reagent used was polyethylenimine (Polysciences) (Chai et al., 2017; Wang et al., 2023; Wei et al., 2024). The weight ratio of the GluA, TARP γ-2, and ABHD6 plasmids was 2:3:0.3125 (the GluA plasmid was 0.8 μg/well). Transfection was terminated 3 h after initiation. HEK 293T cells were dissociated with 0.05% trypsin and then placed on poly-D-lysine-coated coverslips. To prevent cell toxicity due to AMPARs transfection, 200 μM NBQX (Abcam) was added to the culture medium after transfection. The transfected cells were grown for more than 18 h before electrophysiology recording.

### Mice Hippocampal Neuron culture

Place coverslips into 48-well plates, place 100 μL Poly-D-Lysine onto each coverslip, and leave in the incubator for at least 1 hour. Then wash the coverslips with ddH_2_O 3 times, put the coverslips in the hood, and open the covers. Place dissecting instruments into ethanol and then air dry. Put the dissecting pan on ice. Take 5 mL HBS, add 5 µL CaCl_2_ (1M) + 5 µL EDTA (0.5M) + 80 µL Papain (LS003126, Worthington), and then put the fluid mixture in the incubator. Warm plating medium. Dissect the hippocampus in ice-cold Hanks balanced salts solution (HBSS), put in 15mL tube, then transfer it to a culture hood. Wash once with HBS. Clean the enzyme solution, then filter the enzyme solution into the tube. Incubate at 37°C for 20 min. Then remove the enzyme solution and wash with 10mL HF (the HBS with 10% FBS) twice. Aspirate the supernatant from the tissue, and put 720 µL of plating medium. Break down the tissue by trituration using a 1 mL pipette tip 3 times. Pass the cell suspension through a 40µm Nylon cell strainer. Centrifuge the cell mixture at 200× g for 5 min. Remove the mixture and add 2mL of plating medium to cells. After cell counting, ensure that 70,000 cells are planted in each coverslip. Slowly place the cells in the cell incubator for 30 min. Then slowly add 300 µL of plating medium in each well. Finally, the cells are placed in the cell culture incubator. Pipette out 150 µL of media from each well and replace with 180 µL of growth solution (0 Ara-C) at Day 1. Pipette out 150 µL of media from each well and replace with 180 µL of growth solution (4 Ara-C) at Day 4. Pipette out 150 µL of media from each well and replace with 180 µL of growth solution (4 Ara-C) at Day 8. Electrophysiological recordings were performed at Day 14.

### Outside-out patch recording

Recordings were obtained using an EPC 10 patch clamp (HEKA). The data were digitized at 10 kHz. Unless otherwise stated, the holding potential was −60 mV, and series resistance compensation was set to 60–80%, and recordings with series resistances >20 MΩ or the I_leak_ >-200 pA were rejected. The cells were continuously perfused with an external solution containing 150 mM NaCl, 4 mM KCl, 2 mM CaCl_2_,1 mM MgCl_2_, 10 mM HEPES, and 10 mM glucose at pH 7.4 and Osm 315.

Electrodes with a resistance of 4–6 MΩ were pulled from borosilicate glass capillaries (WPI). The internal solution contained 125 mM KF, 33 mM KOH, 2 mM MgCl_2_, 1 mM CaCl_2_, 11 mM EGTA, and 10 mM HEPES at pH 7.4 and Osm 310, and 50 μM spermine was added when indicated. For recording primary hippocampus neurons, 100 μM PTX, 50 μ D-APV, 0.5 μM TTX, and 10 μM NBQX were added into the external solution. Solutions containing L-glutamate (Sigma-Aldrich) were applied by patch membrane perfusion with θ tubes driven by a piezo manipulator (HVA). Glutamate (10 mM) pulses of 1 or 100 ms were applied every 5 s. To calculate the weighted time constant, the time constant of deactivation (τ_w, deact_) and the time constant of desensitization (τ_w, des_) were fitted according to a double exponential function. For AMPAR recovery from desensitization, two-pulse (10 mM glutamate; 100 ms) stimulation was performed at intervals of 1–601 ms. To allow full recovery from desensitization, the sweeps were separated by 5 s. The current evoked by the preceding stimulus was defined as 1, while the other was defined as 2. The recovery ratio was calculated as follows:

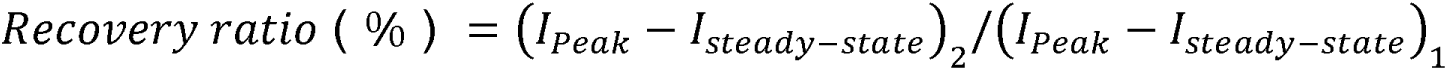

The time constant of recovery from desensitization (τ_rec_) was calculated using nonlinear one-phase exponential decay fitting.

### Affinity Chromatography Experiments and Western Blotting

2.5 μg ABHD6 and 7.5 μg GluA1i (or TARP γ-2) were transfected into a 60-mm dish, a pCAG empty vector was used as a control. After 48 hours, the cells were harvested with PBS, washed once, kept at-80 ℃ overnight, and thawed at 37°C for 1 min. Then the cells were collected with phosphate-buffered saline (PBS) and centrifuged at 13,000× g for 5 min at 4° C to obtain the cell pellets. 300 μL of buffer A (150 mM NaCl, 20 mM HEPES, 2 mM CaCl_2_, 2 mM MgCl_2_, 0.1 mM EDTA, 1% Triton, and protease inhibitors) was added to the cell pellets. Proteins were solubilized by gentle rocking at 4°C for 2 h. Next, the insoluble fractions were removed by centrifugation at 13,000× g for 30 min. A total of 250 μL of supernatant was used for an affinity chromatography assay, and 18 μL were used as inputs. 3 μL anti-Myc Antibody (M20002, Abmart) or anti-Myc Antibody (AE070, ABclonal) with 24 μL of protein G beads were added to samples and rotated overnight at 4 °C. Then, the beads were washed five times with wash buffer (150 mM NaCl, 20 mM HEPES, 2 mM CaCl_2_, 2 mM MgCl_2_, 0.1 mM EDTA, 1% Triton, and protease inhibitors), boiled in SDS sample buffer, and subjected to sodium dodecyl sulfate polyacrylamide gel (SDS-PAGE) electrophoresis.

### Immunostaining Analyses

The coverslips with transfected HEK293T cells were washed once with PBS. Fixed with 4% formaldehyde in PBS for 15 min at room temperature (RT), then permeabilized with 0.2% Triton X-100 in PBS for 15 min at RT, removed the PBS and then blocked with PBS containing 2% goat serum for 30 min at RT. The cells were then incubated for 2 h at RT with the primary antibody (Anti-Glutamate receptor 1 Antibody, AB1504, Millipore; anti-Myc Antibody, M20002, Abmart) diluted in blocking solution. After being washed 3 times every 5 min, the cells were incubated for 30 min at RT with the second antibody (Goat anti-Rabbit IgG (H+L) Highly Cross-Adsorbed Secondary Antibody, Alexa Fluor™ Plus 647, Invitrogen, A32733; Goat anti-Mouse IgG (H+L) Highly Cross-Adsorbed Secondary Antibody, Alexa Fluor ™ 555, A-21424) and washed three to five times with PBS. Images were acquired with a laser scanning confocal microscope (Olympus) and were further analyzed in a blinded fashion using the National Institutes of Health (NIH) ImageJ program (ImageJ, RRID:SCR_003070).

## Statistical analysis

Data were first assessed for normality using the D’Agostino–Pearson test (n<50) or the Kolmogorov-Smirnov test (n> 50), and for equality of variances using the Brown-Forsythe ANOVA test. Depending on the outcome of these tests, data were analyzed by parametric (one-way ANOVA) or non-parametric methods (Kruskal-Wallis test) followed by Tukey’s Honest Significant Difference (HSD) test as a post hoc analysis to determine specific differences among groups. Correlation was evaluated with Pearson correlation analysis. Values of P < 0.05 were considered statistically significant.

## Acknowledgments

This work was supported by grants from the National Key Research and Development Program of China (2024YFF0728700, 2023YFF0724802); R&D Program of Beijing Municipal Commission of Education (KZ20231002529); National Natural Science Foundation of China (32450728, 92468302); Beijing Natural Science Foundation (F251014);Chinese Institutes for Medical Research, Beijing (CX23YZ03,CX24PY01).

## Author contributions

R.C., H.L., H.Y., J.G., L.Y., M.W., and C.Z. designed the research; R.C., H.L., H.Y., Q.L., J.G., S.W. and M.W. conducted the research and analyzed the data; X.G., T.S., Y.Z., D.W. and X.C. analyzed the data; R.C., H.L., H.Y., M.W. and C.Z. wrote the paper.

## Competing interests

The authors declare that they have no conflicts of interest.

---

**Figure EV1.**
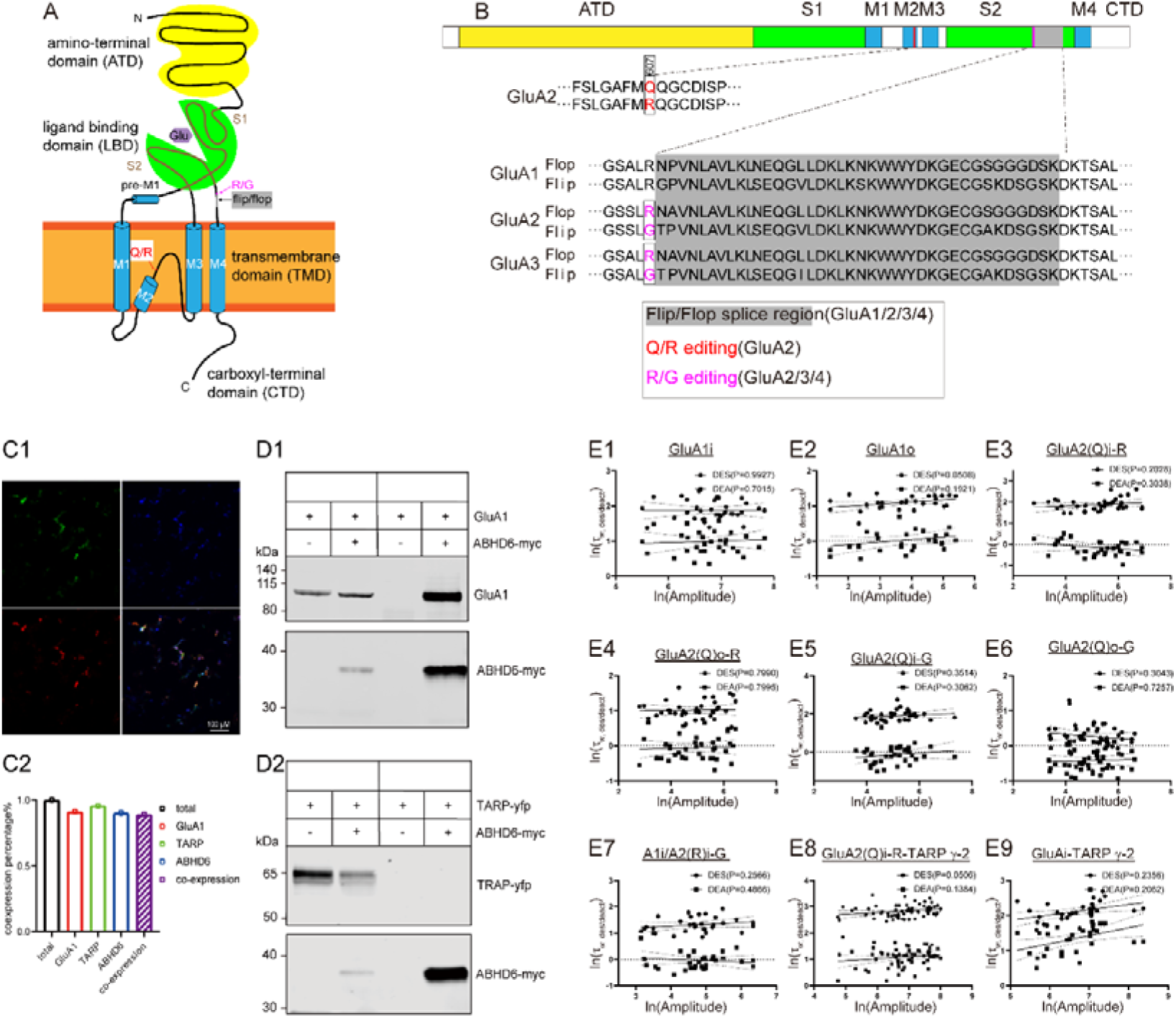
Schematic illustration of the AMPAR subunit; the sequence alignment of RNA splice variants and editing of AMPAR; co-expression validation and Pearson’s correlation between natural logarithm peak amplitude (pA) and the. τ **_w, deact_ and** τ **_w, des_ co-expression with various GluA subunits.** (A) Topology of a single AMPAR subunit in the plasma membrane. Each subunit consists of an extracellular N-terminal domain (NTD), a ligand-binding domain (LBD), a transmembrane domain (TMD, M1–4), and an intracellular C-terminal domain (CTD); the flip/flop splice variants and the RNA editing sites (Q/R and R/G) are also shown. (B) Sequence alignment of RNA splice variants and editing. Q/R editing sites (red letters) (GluA2), R/G editing sites (purple letters) (GluA2, A3), flip/flop splice variants (grey box) (GluA1–A3). Complete sequences can be found in UniProt. Sequences are homologous and conserved in mouse, rat, and human. (C1) Representative confocal images of HEK 293T cells co-transfected with GluA1-flip, ABHD6, and TARP γ-2 plasmids. GluA1-flip: rabbit anti-GluR1 polyclonal antibody (Millipore, Cat. #AB1504, 1:1000) followed by Goat anti-Rabbit IgG (H+L) Highly Cross-Adsorbed Secondary Antibody, Alexa Fluor™ Plus 647 (Invitrogen, Cat. #A32733, 1:2000). ABHD6: detected via its Myc tag; primary antibody Myc-Tag (19C2) mAb (Abmart, Cat. #M20002, 1:1000) and secondary Goat anti-Mouse IgG (H+L) Highly Cross-Adsorbed Secondary Antibody, Alexa Fluor™ 555 (Invitrogen, Cat. #A-21424, 1:2000). (C2) The summary bar graphs of GluA1, TARP γ-2, ABHD6, and co-expression. (D) Affinity chromatography for ABHD6 and TARP γ-2; ABHD6 and GluA1i (or TARP γ-2) were transfected into HEK 293T cells and anti-Myc Antibody (M20002, Abmart) or anti-Myc Antibody (AE070, ABclonal) with 24 μL of protein G beads was added to samples, and then the beads were washed. (E1-E9) Pearson’s correlation and Local Polynomial Regression (loess) with 95% confidence intervals between natural logarithm peak amplitude (pA) and the τ w, deact (ms) of Glutamate (10 mM Glu, 1 ms) induced currents and τ w, des (ms) of Glutamate (10 mM Glu, 100 ms) induced currents in the outside-out patch from HEK 293T cells transfected with various GluA subunits.

**Figure EV2.**
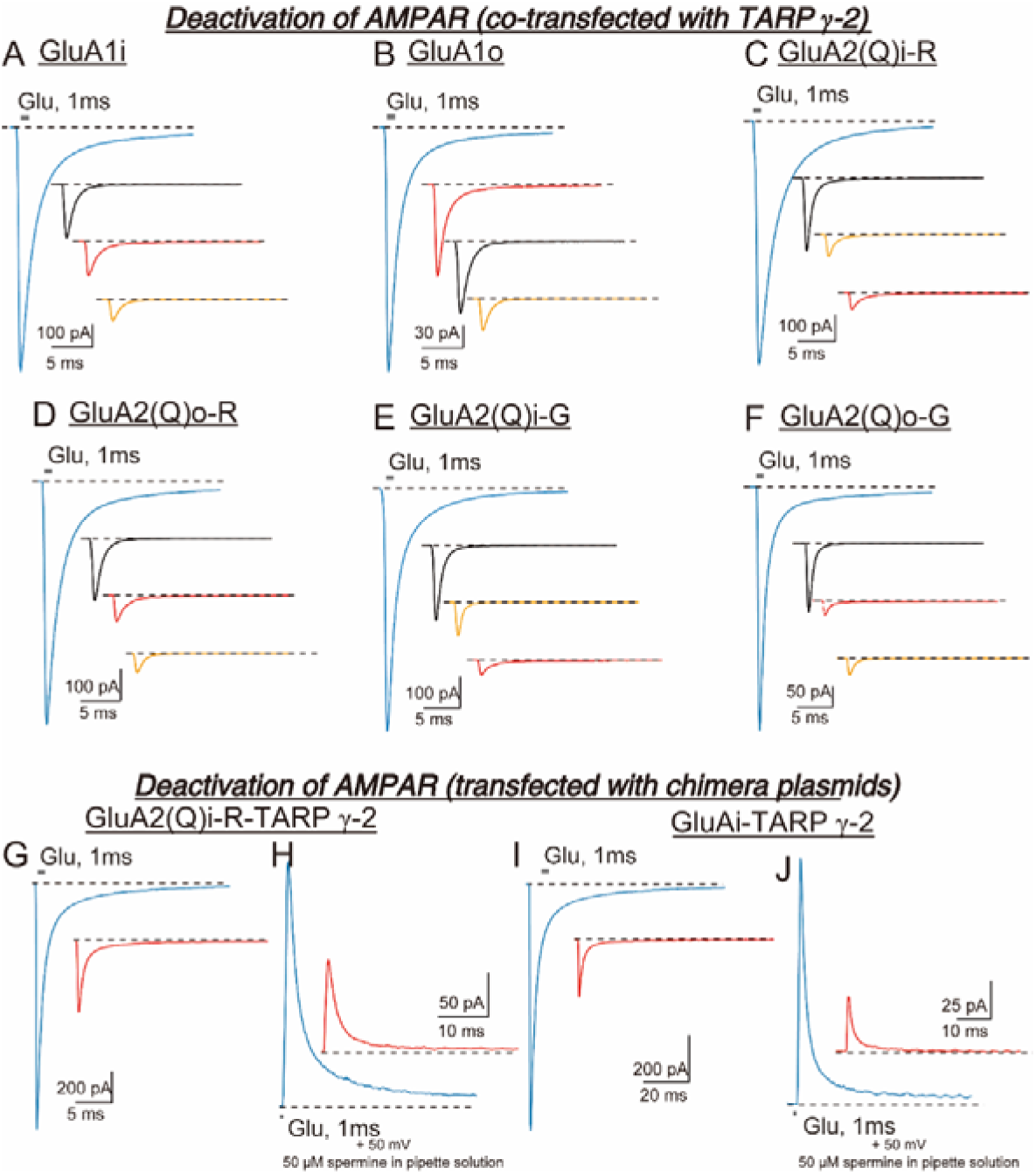
Average traces of deactivation of AMPAR with overexpression of ABHD6 in HEK 293T cells. The average traces of Glutamate (10 mM Glu, 1 ms) induced currents in the outside-out patch from HEK 293T cells transfected with GluA (black), GluA + ABHD6 (orange), GluA + TARP γ-2 (blue), and GluA + TARP γ-2 + ABHD6 (red). (A) GluA1i. (B) GluA1o. (C) GluA2(Q)i-R. (D) GluA2(Q)o-R. (E) GluA2(Q)i-G. (F) GluA2(Q)o-G. (G, H) GluA2(Q)i-R-TARP γ-2 tandem. (I, J) GluA1i-TARP γ-2 tandem.

**Figure EV3.**
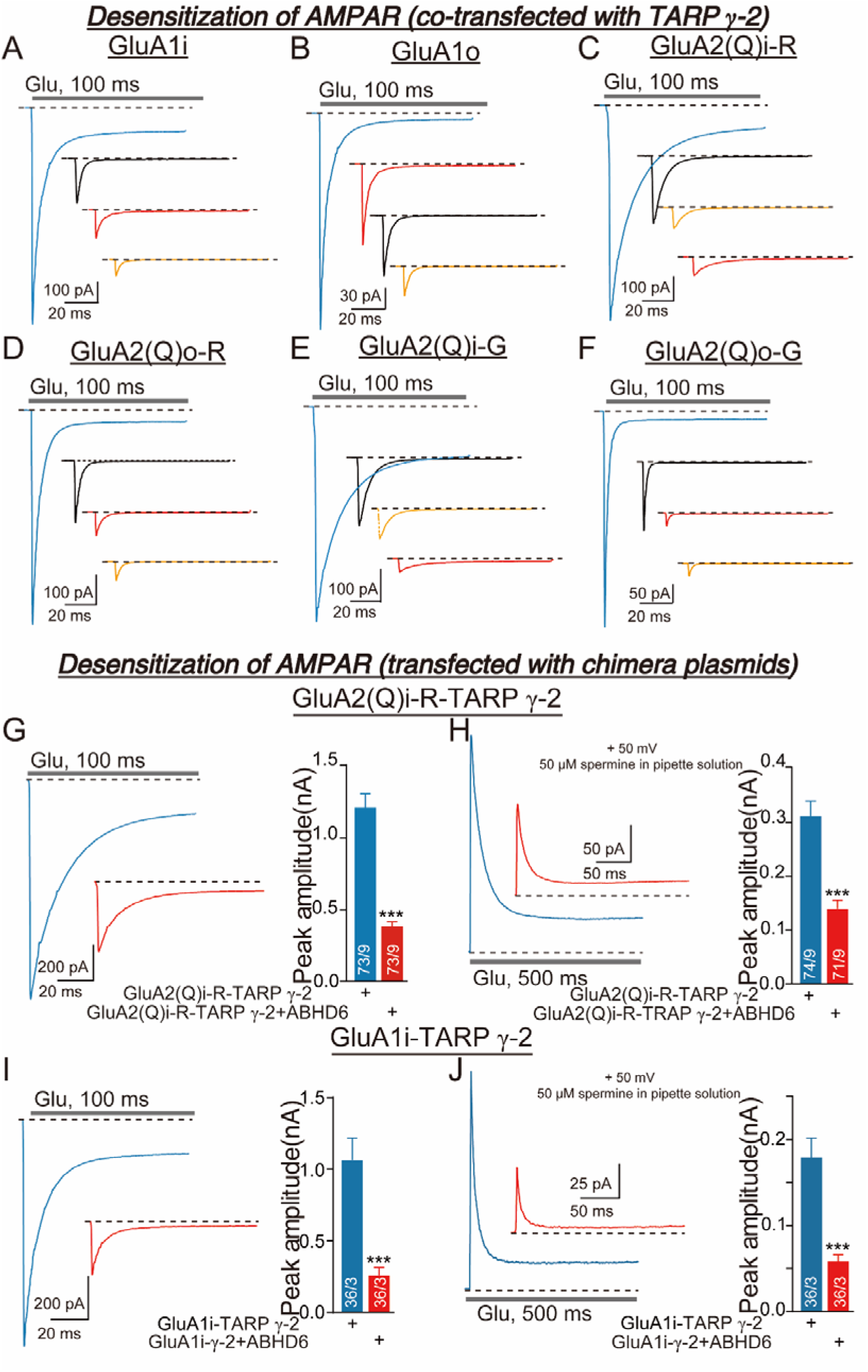
Average traces of desensitization of AMPAR with overexpression of ABHD6 in HEK 293T cells. The average traces of Glutamate (10 mM Glu, 500 ms) induced currents in the outside-out patch from HEK 293T cells transfected with GluA (black), GluA + ABHD6 (orange), GluA + TARP γ-2 (blue), and GluA + TARP γ-2 + ABHD6 (red). (A) GluA1i. (B) GluA1o. (C) GluA2(Q)i-R. (D) GluA2(Q)o-R. (E) GluA2(Q)i-G. (F) GluA2(Q)o-G. (G, H) GluA2(Q)i-R-TARP γ-2 tandem. (I, J) GluA1i-TARP γ-2 tandem.

**Figure EV4.**
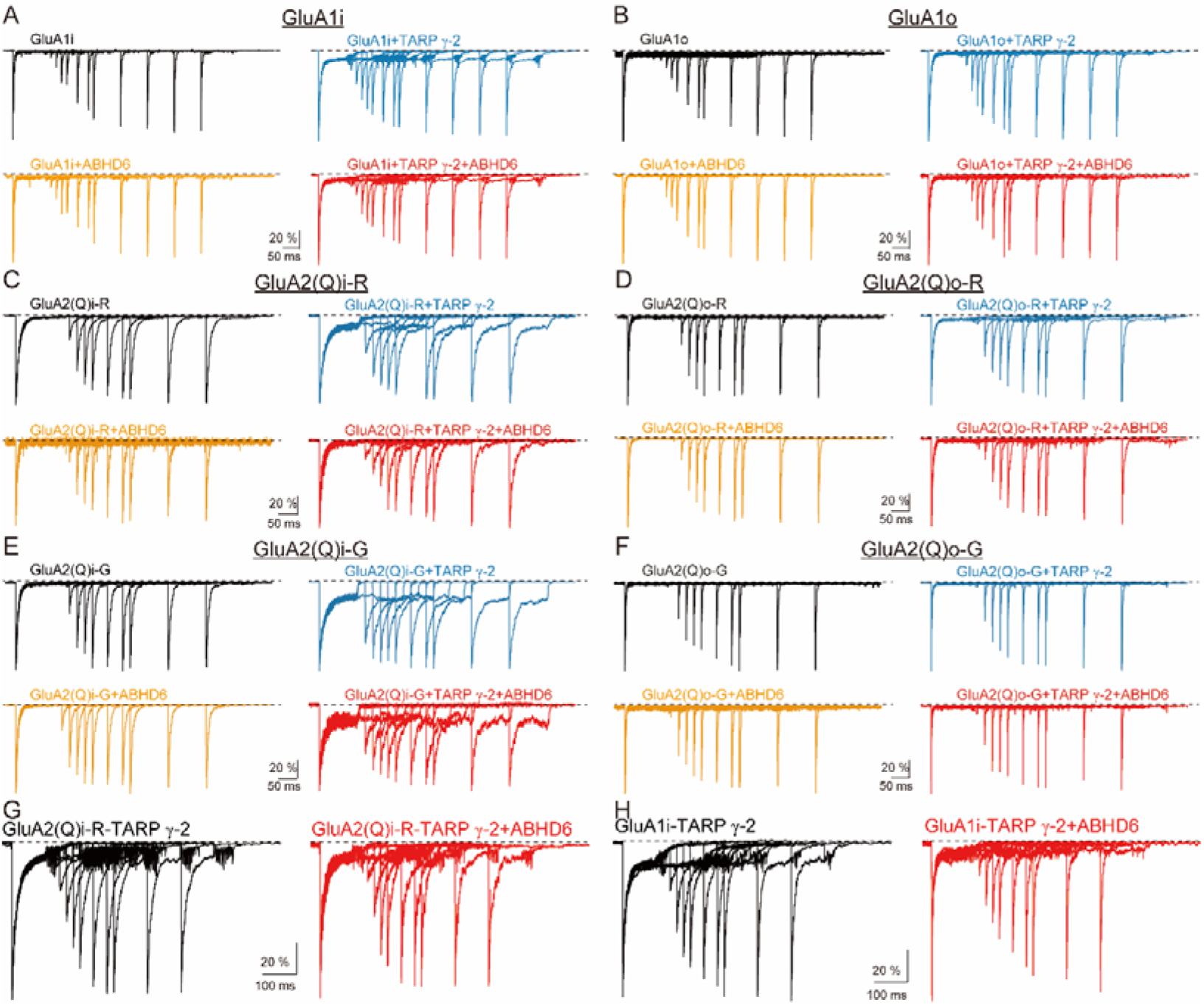
Typical traces of the recovery from desensitization of AMPAR in HEK 293T cells. (A-F) Glutamate (Glu, 10 mM) induced currents in an outside-out patch excised from an HEK 293T cell transfected with GluA (black), GluA + ABHD6 (orange), GluA + TARP γ-2 (blue), and GluA+ TARP γ-2 + ABHD6 (red). The first application of 100 ms glutamate was followed by a second glutamate application at increasing pulse intervals at-60 mV. The typical traces from a cell are normalized and aligned to the peak. The typical traces of the recovery from desensitization. (A) GluA1i. (B) GluA1o. (C) GluA2(Q)i-R. (D) GluA2(Q)o-R. (E) GluA2(Q)i-G. (F) GluA2(Q)o-G. (G) GluA2(Q)i-R-TARP γ-2 tandem. (H) GluA1i-TARP γ-2 tandem.

**Figure EV5.**
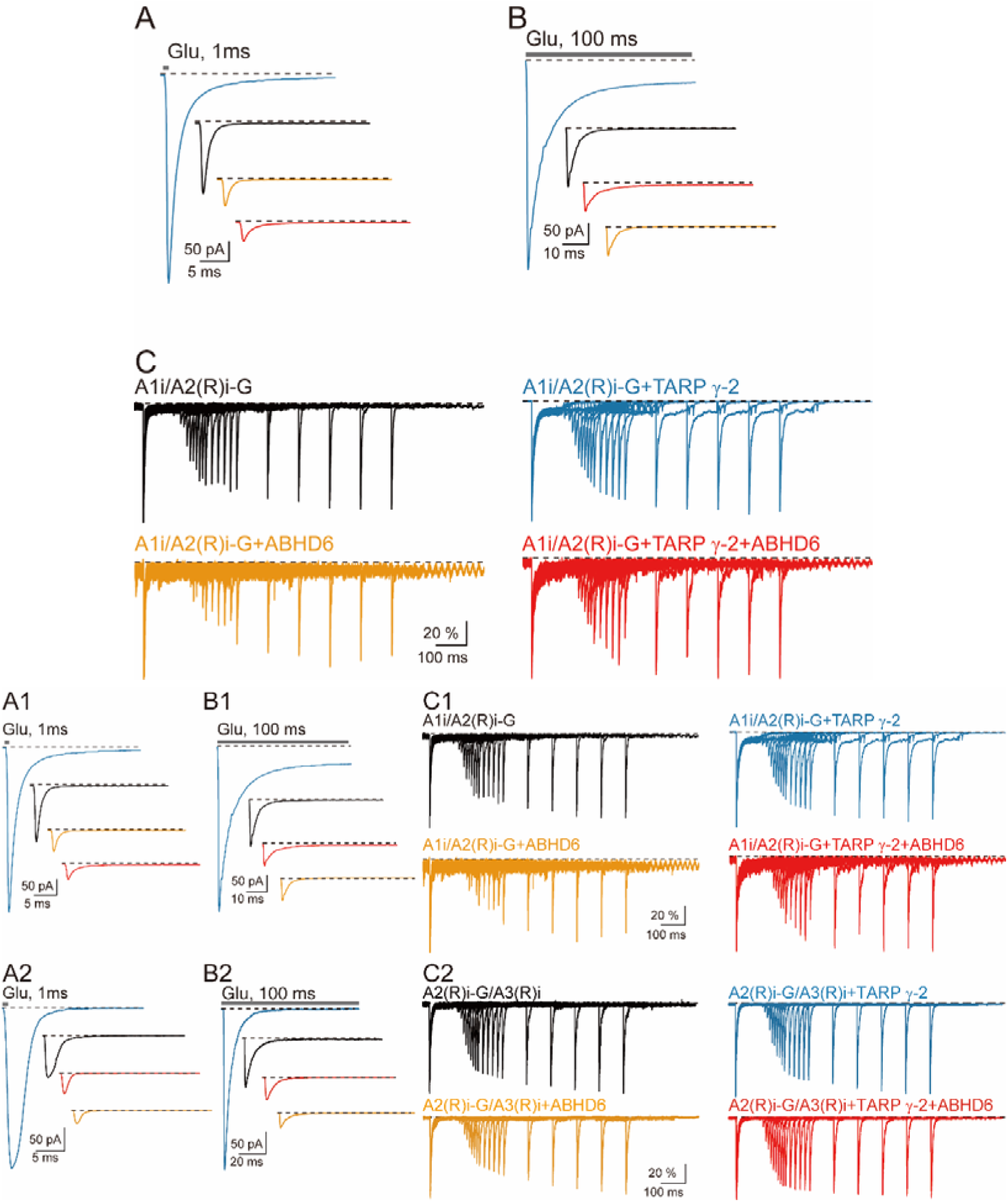
Average traces of the deactivation, desensitization, and recovery from desensitization of GluA1i/GluA2(R)i-G (or GluA2(R)i-G/GluA3(R)i) receptors-TARP. γ**-2 complexes in HEK 293T cells.** (A) The average traces of the τ _w, deact_ of glutamate (10 mM Glu, 1 ms) induced currents in the outside-out patch recorded at-60 mV from HEK 293T cells transfected with GluA1i/GluA2(R)i-G or GluA2(R)i-G/GluA3(R)i (black), GluA1i/GluA2(R)i-G or GluA2(R)i-G/GluA3(R)i + ABHD6 (orange), GluA1i/GluA2(R)i-G or GluA2(R)i-G/GluA3(R)i + TARP γ-2 (blue), and GluA1i/GluA2(R)i-G or GluA2(R)i-G/GluA3(R)i + TARP γ-2 + ABHD6 (red). (B) The average traces of the τ _w, des_, and peak amplitude of glutamate (10 mM Glu, 100 ms) induced currents in the outside-out patch recorded at-60 mV from HEK 293T cells transfected with GluA1i/GluA2(R)i-G or GluA2(R)i-G/GluA3(R)i (black), GluA1i/GluA2(R)i-G or GluA2(R)i-G/GluA3(R)i + ABHD6 (orange), GluA1i/GluA2(R)i-G or GluA2(R)i-G/GluA3(R)i + TARP γ-2 (blue), and GluA1i/GluA2(R)i-G or GluA2(R)i-G/GluA3(R)i + TARP γ-2 + ABHD6 (red). (C) The typical trace of recovery from desensitization. Glutamate (Glu, 10 mM) induced currents in an outside-out patch excised from an HEK 293T cell transfected with GluA1i/GluA2(R)i-G or GluA2(R)i-G/GluA3(R)i (black), GluA1i/GluA2(R)i-G or GluA2(R)i-G/GluA3(R)i + ABHD6 (orange), GluA1i/GluA2(R)i-G or GluA2(R)i-G/GluA3(R)i + TARP γ-2 (blue), and GluA1i/GluA2(R)i-G or GluA2(R)i-G/GluA3(R)i + TARP γ-2 + ABHD6 (red). The first application of 100 ms glutamate was followed by a second glutamate application at increasing pulse intervals at-60 mV.

**Figure EV7.**
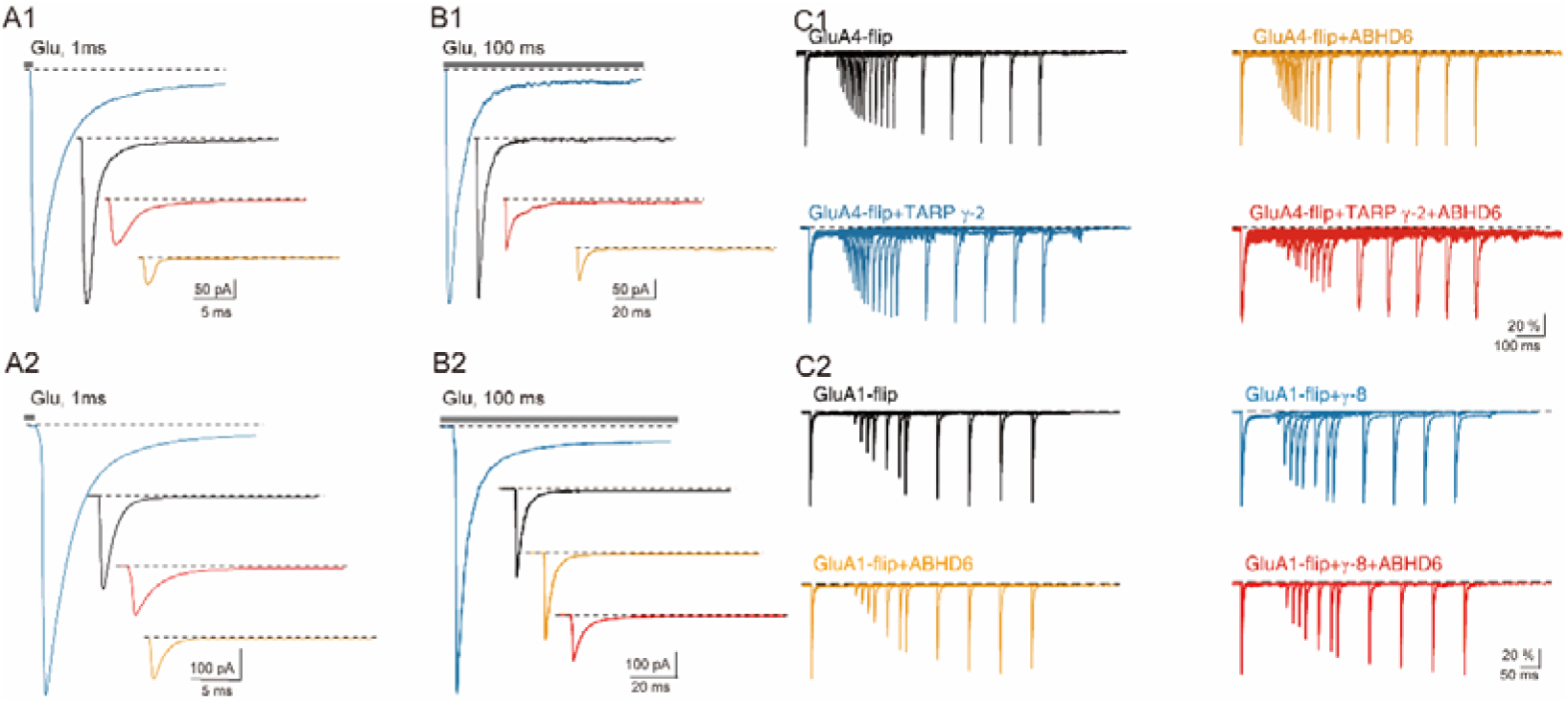
Average traces of the deactivation, desensitization, and recovery from desensitization of GluA4i-TARP. γ**-2 complexes and GluA1i-TARP** γ**-8 complexes in HEK 293T cells.** (A) The average traces of the τ _w, deact_ of glutamate (10 mM Glu, 1 ms) induced currents in the outside-out patch recorded at-60 mV from HEK 293T cells transfected with GluA4i or GluA1i (black), GluA4i or GluA1i (black) + ABHD6 (orange), GluA4i or GluA1i + TARP γ-2 (blue), or GluA4i or GluA1i + TARP γ-2 + ABHD6 (red). (B) The average traces of the τ _w, des_, and peak amplitude of glutamate (10 mM Glu, 100 ms) induced currents in the outside-out patch recorded at-60 mV from HEK 293T cells transfected with GluA4i or GluA1i (black), GluA4i or GluA1i (black) + ABHD6 (orange), GluA4i or GluA1i + TARP γ-2 (blue), or GluA4i or GluA1i + TARP γ-2 + ABHD6 (red). (C) The typical trace of recovery from desensitization. glutamate (Glu, 10 mM) induced currents in an outside-out patch excised from an HEK 293T cell transfected with GluA4i or GluA1i (black), GluA4i or GluA1i (black) + ABHD6 (orange), GluA4i or GluA1i + TARP γ-2 (blue), or GluA4i or GluA1i + TARP γ-2 + ABHD6 (red). The first application of 100 ms glutamate was followed by a second glutamate application at increasing pulse intervals at-60 mV.

**Table EV1.1.**
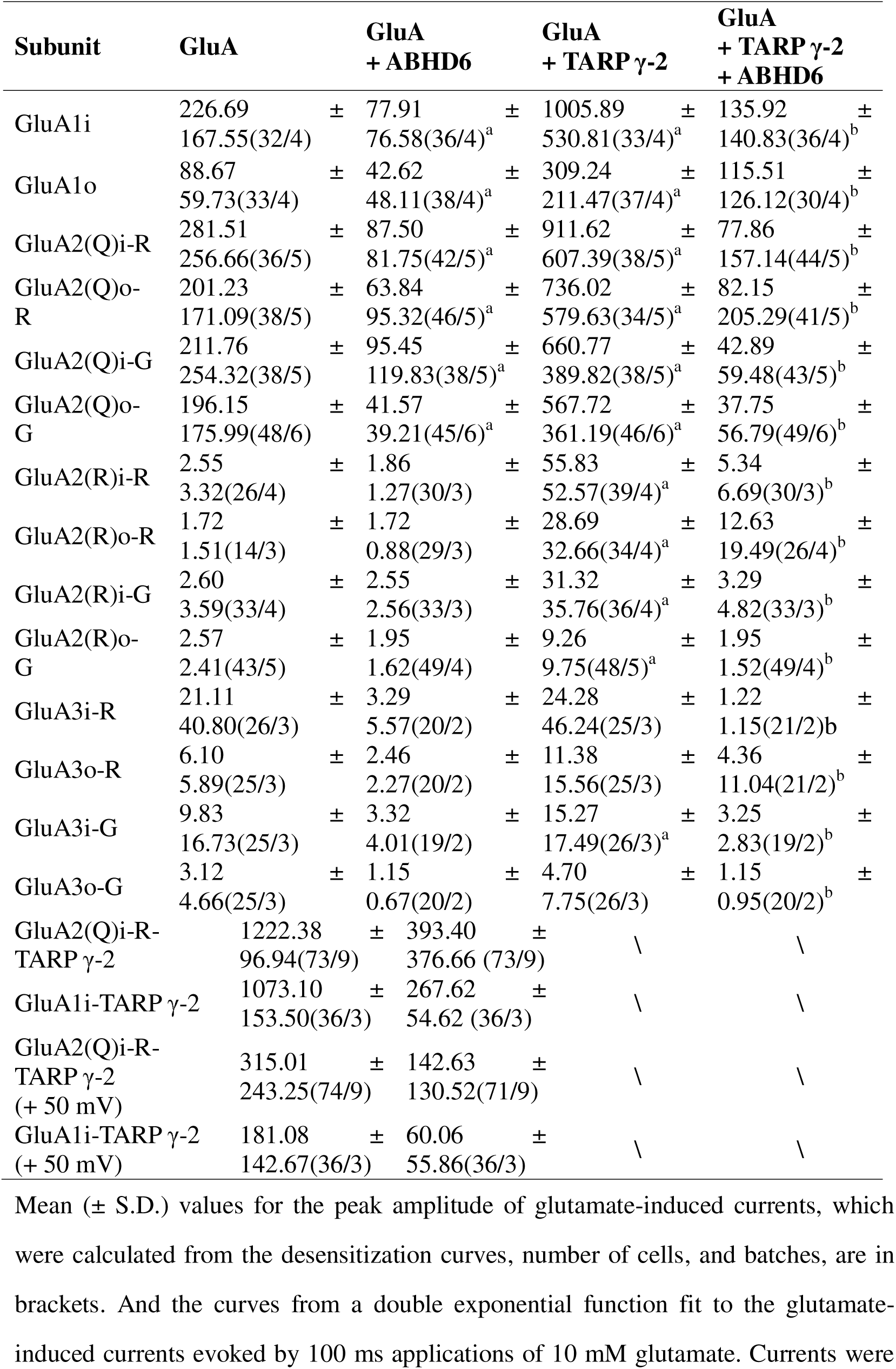
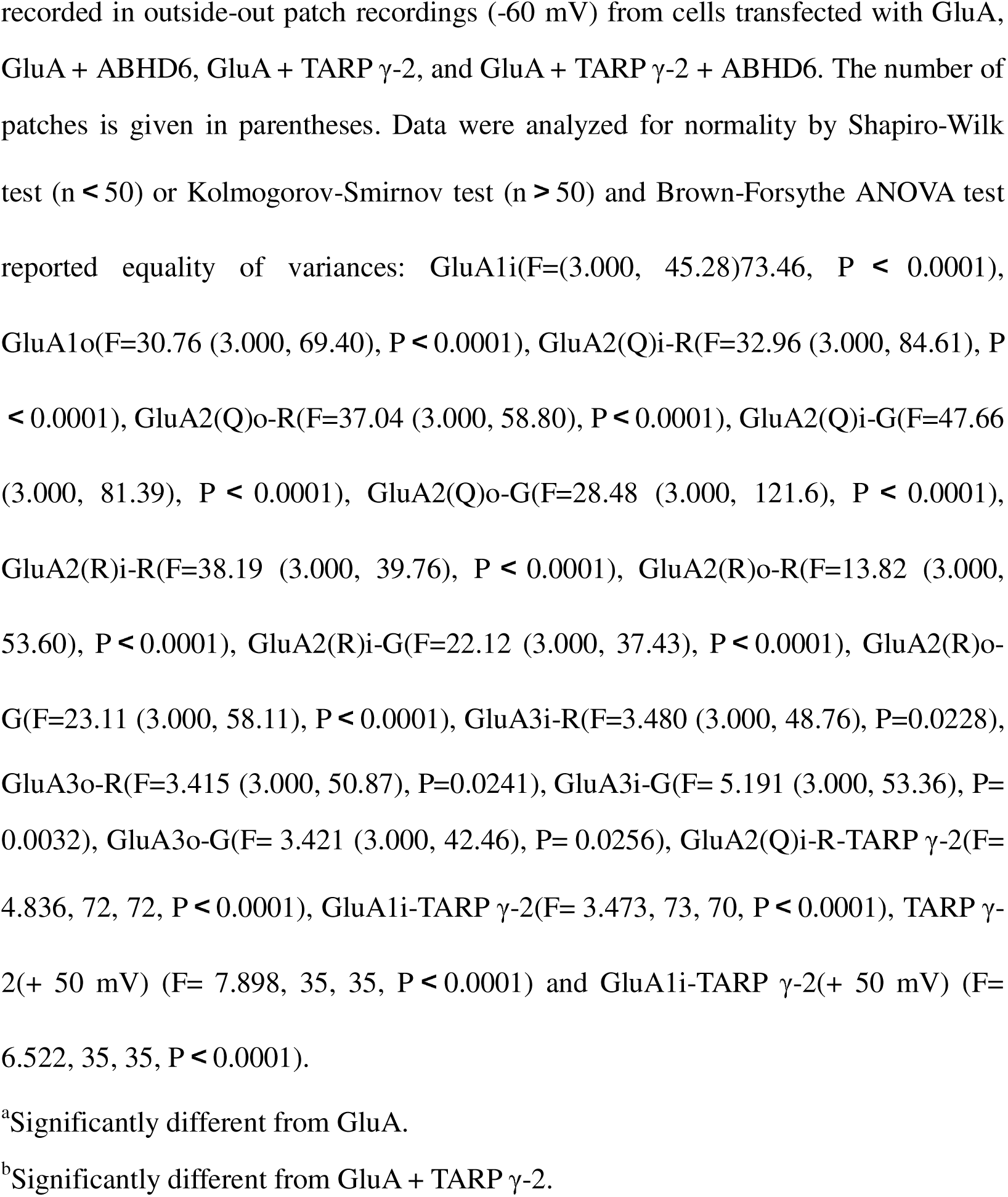
Summary of peak amplitude (pA) of GluAs when co-transfected with/without γ-2 or/and ABHD6.

| Subunit | GluA | | GluA<br>+ ABHD6 | | GluA<br>+ TARP γ-2 | | GluA<br>+ TARP γ-2<br>+ ABHD6 |
| --- | --- | --- | --- | --- | --- | --- | --- |
| GluA1i | 226.69<br>167.55(32/4) | ± | 77.91<br>76.58(36/4)<sup>a</sup> | ± | 1005.89<br>530.81(33/4)<sup>a</sup> | ± | 135.92<br>140.83(36/4)<sup>b</sup> |
| GluA1o | 88.67<br>59.73(33/4) | ± | 42.62<br>48.11(38/4)<sup>a</sup> | ± | 309.24<br>211.47(37/4)<sup>a</sup> | ± | 115.51<br>126.12(30/4)<sup>b</sup> |
| GluA2(Q)i-R | 281.51<br>256.66(36/5) | ± | 87.50<br>81.75(42/5)<sup>a</sup> | ± | 911.62<br>607.39(38/5)<sup>a</sup> | ± | 77.86<br>157.14(44/5)<sup>b</sup> |
| GluA2(Q)o-R | 201.23<br>171.09(38/5) | ± | 63.84<br>95.32(46/5)<sup>a</sup> | ± | 736.02<br>579.63(34/5)<sup>a</sup> | ± | 82.15<br>205.29(41/5)<sup>b</sup> |
| GluA2(Q)i-G | 211.76<br>254.32(38/5) | ± | 95.45<br>119.83(38/5)<sup>a</sup> | ± | 660.77<br>389.82(38/5)<sup>a</sup> | ± | 42.89<br>59.48(43/5)<sup>b</sup> |
| GluA2(Q)o-G | 196.15<br>175.99(48/6) | ± | 41.57<br>39.21(45/6)<sup>a</sup> | ± | 567.72<br>361.19(46/6)<sup>a</sup> | ± | 37.75<br>56.79(49/6)<sup>b</sup> |
| GluA2(R)i-R | 2.55<br>3.32(26/4) | ± | 1.86<br>1.27(30/3) | ± | 55.83<br>52.57(39/4)<sup>a</sup> | ± | 5.34<br>6.69(30/3)<sup>b</sup> |
| GluA2(R)o-R | 1.72<br>1.51(14/3) | ± | 1.72<br>0.88(29/3) | ± | 28.69<br>32.66(34/4)<sup>a</sup> | ± | 12.63<br>19.49(26/4)<sup>b</sup> |
| GluA2(R)i-G | 2.60<br>3.59(33/4) | ± | 2.55<br>2.56(33/3) | ± | 31.32<br>35.76(36/4)<sup>a</sup> | ± | 3.29<br>4.82(33/3)<sup>b</sup> |
| GluA2(R)o-G | 2.57<br>2.41(43/5) | ± | 1.95<br>1.62(49/4) | ± | 9.26<br>9.75(48/5)<sup>a</sup> | ± | 1.95<br>1.52(49/4)<sup>b</sup> |
| GluA3i-R | 21.11<br>40.80(26/3) | ± | 3.29<br>5.57(20/2) | ± | 24.28<br>46.24(25/3) | ± | 1.22<br>1.15(21/2)<sup>b</sup> |
| GluA3o-R | 6.10<br>5.89(25/3) | ± | 2.46<br>2.27(20/2) | ± | 11.38<br>15.56(25/3) | ± | 4.36<br>11.04(21/2)<sup>b</sup> |
| GluA3i-G | 9.83<br>16.73(25/3) | ± | 3.32<br>4.01(19/2) | ± | 15.27<br>17.49(26/3)<sup>a</sup> | ± | 3.25<br>2.83(19/2)<sup>b</sup> |
| GluA3o-G | 3.12<br>4.66(25/3) | ± | 1.15<br>0.67(20/2) | ± | 4.70<br>7.75(26/3) | ± | 1.15<br>0.95(20/2)<sup>b</sup> |
| GluA2(Q)i-R-TARP γ-2 | 1222.38<br>96.94(73/9) | ± | 393.40<br>376.66 (73/9) | ± | \ | | \ |
| GluA1i-TARP γ-2 | 1073.10<br>153.50(36/3) | ± | 267.62<br>54.62 (36/3) | ± | \ | | \ |
| GluA2(Q)i-R-TARP γ-2<br>(+ 50 mV) | 315.01<br>243.25(74/9) | ± | 142.63<br>130.52(71/9) | ± | \ | | \ |
| GluA1i-TARP γ-2<br>(+ 50 mV) | 181.08<br>142.67(36/3) | ± | 60.06<br>55.86(36/3) | ± | \ | | \ |

Mean ( ± S.D.) values for the peak amplitude of glutamate-induced currents, which were calculated from the desensitization curves, number of cells, and batches, are in brackets. And the curves from a double exponential function fit to the glutamate-induced currents evoked by 100 ms applications of 10 mM glutamate. Currents were
recorded in outside-out patch recordings (-60 mV) from cells transfected with GluA, GluA + ABHD6, GluA + TARP γ-2, and GluA + TARP γ-2 + ABHD6. The number of patches is given in parentheses. Data were analyzed for normality by Shapiro-Wilk test ( $n < 50$ ) or Kolmogorov-Smirnov test ( $n > 50$ ) and Brown-Forsythe ANOVA test reported equality of variances: GluA1i(F=(3.000, 45.28)73.46, $P < 0.0001$ ), GluA1o(F=30.76 (3.000, 69.40), $P < 0.0001$ ), GluA2(Q)i-R(F=32.96 (3.000, 84.61), $P$ $< 0.0001$ ), GluA2(Q)o-R(F=37.04 (3.000, 58.80), $P < 0.0001$ ), GluA2(Q)i-G(F=47.66 (3.000, 81.39), $P < 0.0001$ ), GluA2(Q)o-G(F=28.48 (3.000, 121.6), $P < 0.0001$ ), GluA2(R)i-R(F=38.19 (3.000, 39.76), $P < 0.0001$ ), GluA2(R)o-R(F=13.82 (3.000, 53.60), $P < 0.0001$ ), GluA2(R)i-G(F=22.12 (3.000, 37.43), $P < 0.0001$ ), GluA2(R)o- G(F=23.11 (3.000, 58.11), $P < 0.0001$ ), GluA3i-R(F=3.480 (3.000, 48.76), $P=0.0228$ ), GluA3o-R(F=3.415 (3.000, 50.87), $P=0.0241$ ), GluA3i-G(F= 5.191 (3.000, 53.36), $P=$ 0.0032), GluA3o-G(F= 3.421 (3.000, 42.46), $P= 0.0256$ ), GluA2(Q)i-R-TARP γ-2(F= 4.836, 72, 72, $P < 0.0001$ ), GluA1i-TARP γ-2(F= 3.473, 73, 70, $P < 0.0001$ ), TARP $\gamma$ - 2(+ 50 mV) (F= 7.898, 35, 35, $P < 0.0001$ ) and GluA1i-TARP γ-2(+ 50 mV) (F= 6.522, 35, 35, $P < 0.0001$ ).
<sup>a</sup>Significantly different from GluA.
<sup>b</sup>Significantly different from GluA + TARP γ-2.

**Table EV1.2.**
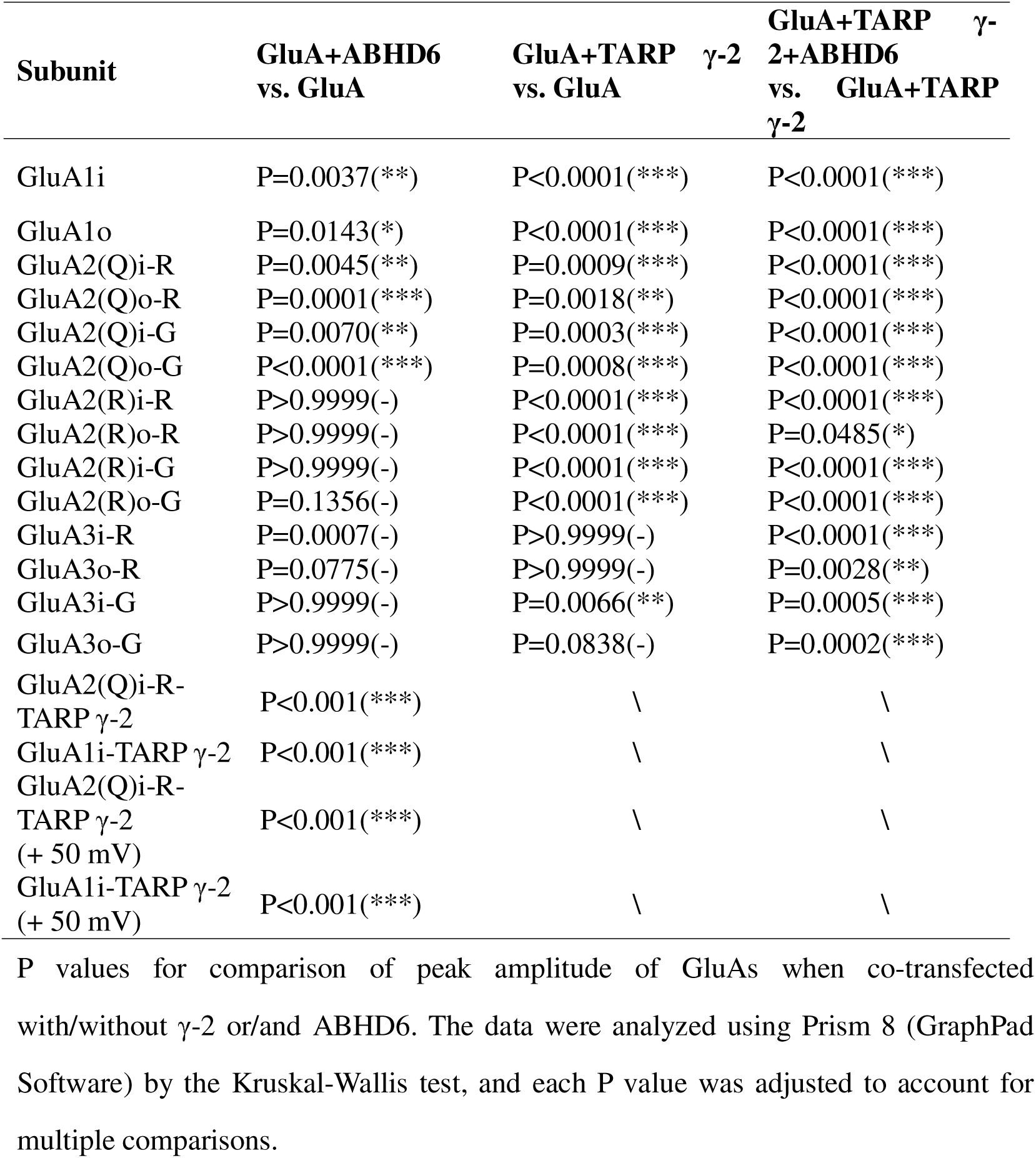
Summary of P values for comparison of peak amplitude.

**Table EV2.1.** Summary of τ _w, deact_ (ms) of GluAs when co-transfected with/without γ-2 or/and ABHD6.

| Subunit | GluA | | GluA<br>+ ABHD6 | | GluA<br>+ TARP γ-2 | | GluA<br>+ TARP γ-2<br>+ ABHD6 | |
| --- | --- | --- | --- | --- | --- | --- | --- | --- |
| GluA1i | 0.99<br>0.21(32/4) | ± | 0.98<br>0.41(36/4) | ± | 2.88<br>1.16(33/4)<sup>a</sup> | ± | 1.38<br>±0.38(36/4)<sup>b</sup> | |
| GluA1o | 1.09<br>0.27(33/4) | ± | 1.00<br>0.35(36/4) | ± | 1.91<br>0.81(37/4)<sup>a</sup> | ± | 1.25<br>0.49(30/4)<sup>b</sup> | ± |
| GluA2(Q)i-R | 1.03<br>1.05(36/5) | ± | 1.15<br>±0.77(43/5) | | 3.68<br>3.13(38/5)<sup>a</sup> | ± | 1.78<br>1.39(43/5)<sup>b</sup> | ± |
| GluA2(Q)o-R | 0.97<br>0.40(38/5) | ± | 1.07<br>0.42(47/5) | ± | 2.78<br>1.43(34/5)<sup>a</sup> | ± | 1.66<br>0.95(46/5)<sup>b</sup> | ± |
| GluA2(Q)i-G | 0.96<br>0.55(37/5) | ± | 0.81<br>0.37(38/5) | ± | 2.44<br>1.46(38/5)<sup>a</sup> | ± | 1.79<br>1.26(44/5) | ± |
| GluA2(Q)o-G | 0.68<br>0.21(48/6) | ± | 0.76<br>0.27(47/6) | ± | 1.90<br>1.30(46/6)<sup>a</sup> | ± | 1.14<br>0.60(50/6)<sup>b</sup> | ± |
| GluA2(Q)i-R-TARP γ-2 | 3.23±<br>1.15(73/9) | | 2.34±<br>1.27(73/9) | | | \ | | \ |
| GluA1i-TARP γ-2 | 4.19±<br>1.69(36/3) | | 2.31±<br>1.11(36/3) | | | \ | | \ |
| GluA2(Q)i-R-TARP γ-2<br>(+ 50 mV) | 2.81<br>1.80(74/9) | ± | 1.71<br>0.93(71/9) | ± | | \ | | \ |
| GluA1i-TARP γ-2<br>(+ 50 mV) | 2.64<br>1.23(36/3) | ± | 1.86<br>1.05(36/3) | ± | | \ | | \ |

Mean ( ± S.D.) values for the time constant of deactivation ( $\tau_{w, deact}$ ). The deactivation curves from a double exponential function fit to the glutamate-induced currents evoked by 1 ms applications of 10 mM glutamate. Currents were recorded in outside-out patch recordings (-60 mV) from cells transfected with GluA, GluA + ABHD6, GluA + TARP γ-2, and GluA + TARP γ-2 + ABHD6. The number of patches is given in parentheses.
<sup>a</sup>Significantly different from GluA.
<sup>b</sup>Significantly different from GluA + TARP γ-2.

**Table EV2.2.**
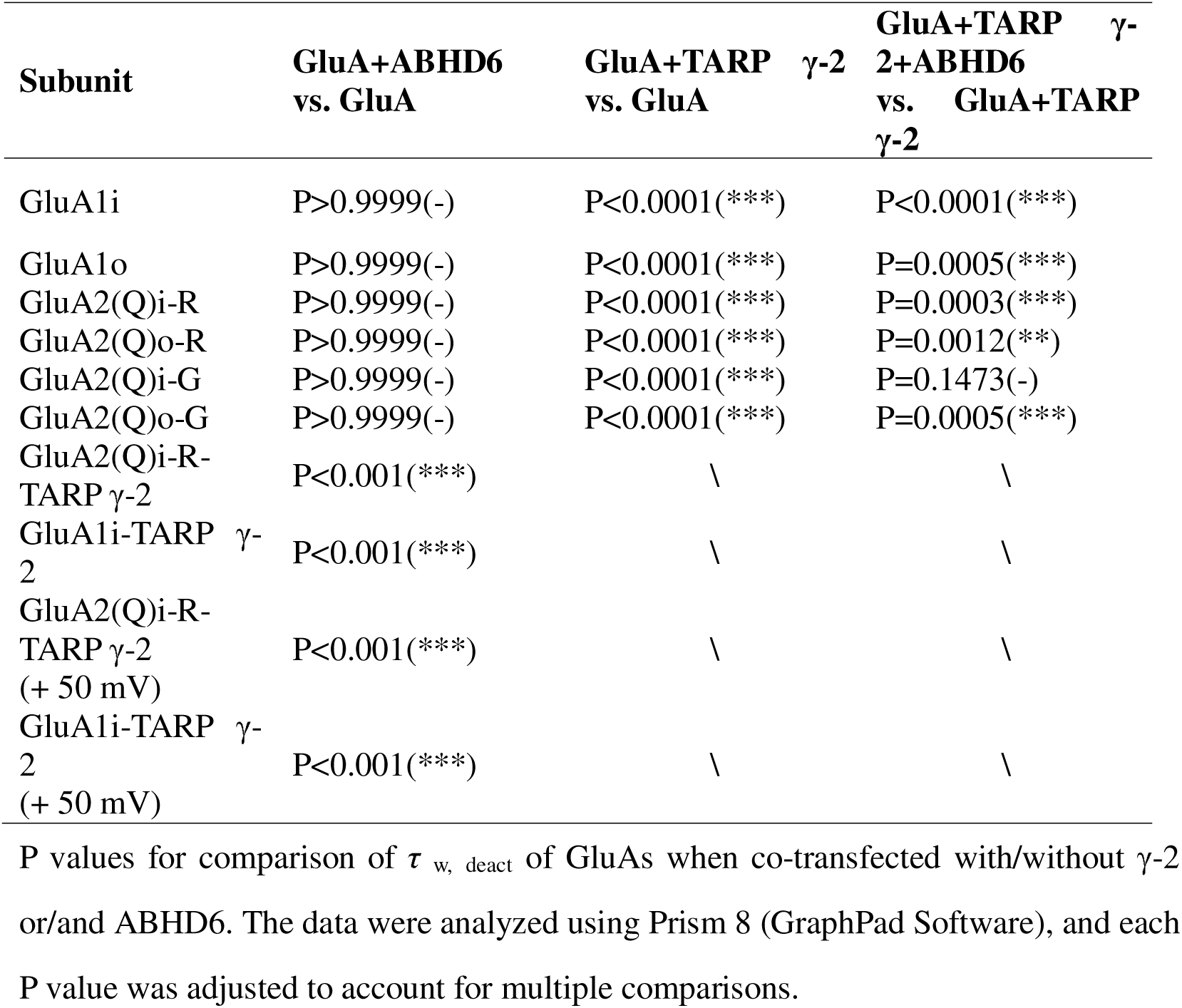
Summary of P values for comparison of τ _w, deact_.

**Table EV3.1.** Summary of τ _w, des_ (ms) of GluAs when co-transfected with/without γ-2 or/and ABHD6.

| Subunit | GluA | | GluA<br>+ ABHD6 | | GluA<br>+ TARP γ-2 | | GluA<br>+ TARP γ-2<br>+ ABHD6 | |
| --- | --- | --- | --- | --- | --- | --- | --- | --- |
| GluA1i | 2.91<br>0.53(32/4) | ± | 2.59<br>0.78(36/4) | ± | 6.72<br>1.86(33/4)<sup>a</sup> | ± | 4.14<br>1.35(36/4)<sup>b</sup> | ± |
| GluA1o | 3.39<br>0.70(33/4) | ± | 2.90<br>0.69(38/4) | ± | 5.52<br>1.59(37/4)<sup>a</sup> | ± | 3.50<br>1.15(30/4)<sup>b</sup> | ± |
| GluA2(Q)i-R | 6.82<br>2.19(36/5) | ± | 6.26<br>1.83(42/5) | ± | 15.43<br>4.87(38/5)<sup>a</sup> | ± | 9.13<br>3.41(44/5)<sup>b</sup> | ± |
| GluA2(Q)o-R | 2.82<br>0.81(38/5) | ± | 2.66<br>0.78(46/5) | ± | 4.71<br>1.48(34/5)<sup>a</sup> | ± | 3.08<br>1.33(41/5)<sup>b</sup> | ± |
| GluA2(Q)i-G | 7.00<br>2.11(38/5) | ± | 6.25<br>2.0(38/5) | ± | 15.85<br>3.95(38/5)<sup>a</sup> | ± | 9.52<br>3.93(43/5)<sup>b</sup> | ± |
| GluA2(Q)o-G | 1.36<br>0.58(48/6) | ± | 1.36<br>0.51(45/6) | ± | 2.01<br>0.60(46/6)<sup>a</sup> | ± | 1.43<br>0.50(49/6)<sup>b</sup> | ± |
| GluA2(Q)i-R-TARP γ-2 | 17.47±<br>4.68(73/9) | | 11.78±<br>3.20(73/9) | | \ | | \ | |
| GluA1i-TARP γ-2 | 8.78±<br>2.76(36/3) | | 5.80±<br>2.12(36/3) | | \ | | \ | |
| GluA2(Q)i-R-TARP γ-2<br>(+ 50 mV) | 15.52±<br>4.48(74/9) | | 10.31<br>3.71(71/9) | ± | \ | | \ | |
| GluA1i-TARP γ-2<br>(+ 50 mV) | 6.81±<br>2.01(36/3) | | 5.64<br>1.85(36/3) | ± | \ | | \ | |

Mean ( ± S.D.) values for the time constant of desensitization ( $\tau_{w, des}$ ) from the desensitization curves. Currents were recorded in outside-out patch recordings (-60 mV) from cells transfected with GluA, GluA + ABHD6, GluA + TARP- γ-2, and GluA + TARP- γ-2 + ABHD6. The number of patches is given in parentheses.
<sup>a</sup>Significantly different from GluA.
<sup>b</sup>Significantly different from GluA + TARP γ-2.

**Table EV3.2.**
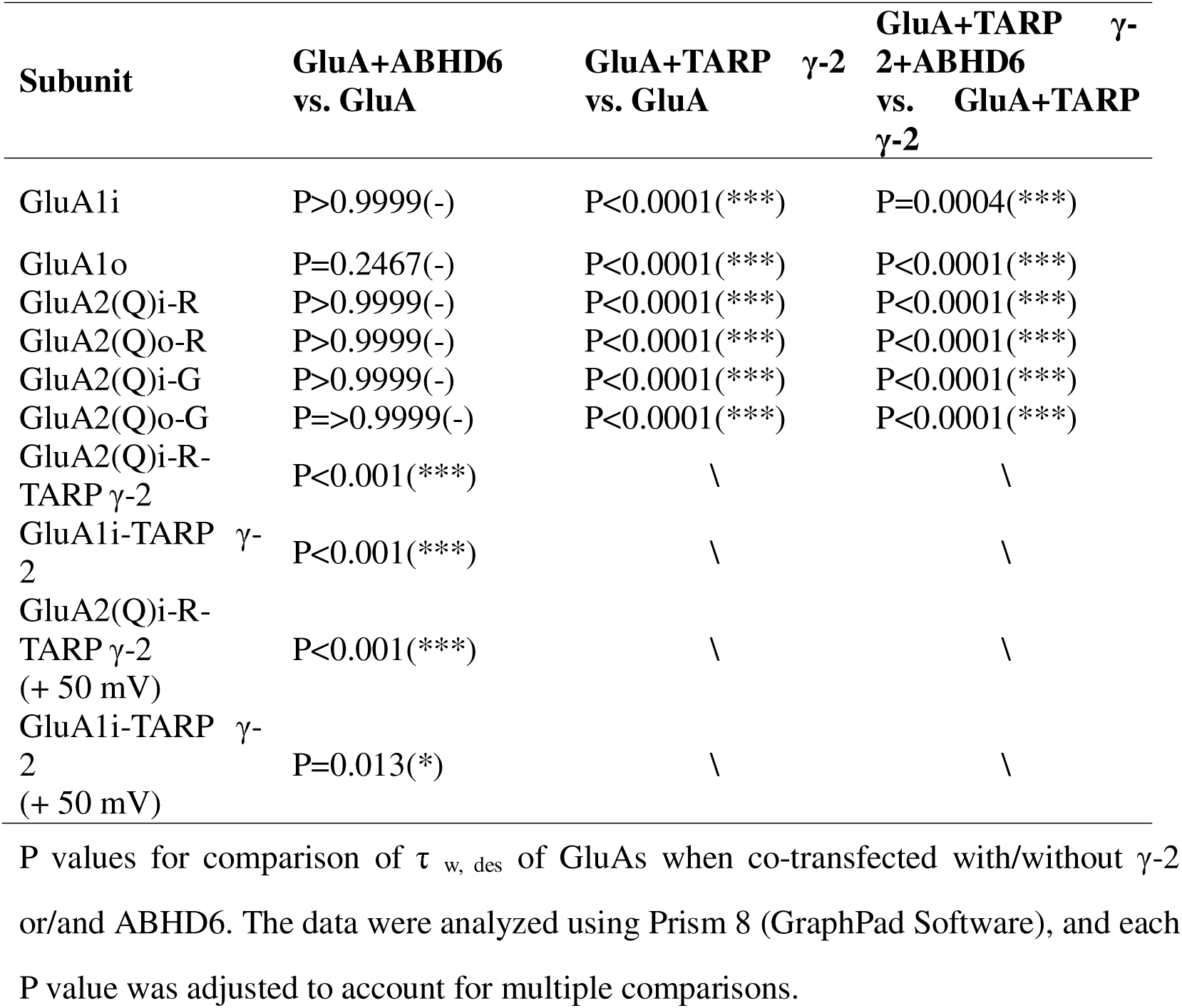
Summary of P values for comparison of τ _w, des_.

**Table EV4.1.** Summary of τ _w, rec_ (ms) of GluAs when co-transfected with/without γ-2 or/and ABHD6.

| Subunit | GluA | | GluA<br>+ ABHD6 | | GluA<br>+ TARP γ-2 | | GluA<br>+ TARP γ-2<br>+ ABHD6 | |
| --- | --- | --- | --- | --- | --- | --- | --- | --- |
| GluA1i | 223.90<br>124.60(32/4) | ± | 225.00<br>201.39(29/4) | ± | 80.66<br>22.56(33/4)<sup>a</sup> | ± | 126.50<br>58.83(30/4)<sup>b</sup> | ± |
| GluA1o | 182.60<br>87.20(28/4) | ± | 165.60<br>39.86(25/4) | ± | 100.70<br>27.76(33/4)<sup>a</sup> | ± | 119.90<br>41.56(26/4) | ± |
| GluA2(Q)i-R | 77.17<br>41.26(36/5) | ± | 75.89<br>32.54(38/5) | ± | 56.98<br>17.86(42/5)<sup>a</sup> | ± | 66.44<br>33.63(27/5) | ± |
| GluA2(Q)o-R | 67.60<br>28.29(35/5) | ± | 68.87<br>36.06(33/5) | ± | 54.05<br>17.56(33/5) | ± | 59.95<br>26.61(33/5) | ± |
| GluA2(Q)i-G | 52.29<br>20.93(37/5) | ± | 56.23<br>37.72(29/5) | ± | 40.85<br>27.86(35/5)<sup>a</sup> | ± | 47.39<br>26.72(31/5) | ± |
| GluA2(Q)o-G | 50.48<br>16.68(44/6) | ± | 58.12<br>33.49(39/6) | ± | 50.39<br>18.40(45/6) | ± | 55.68<br>29.88(35/6) | ± |
| GluA2(Q)i-R-TARP γ-2 | 77.61±<br>20.99(63/9) | | 82.66±<br>20.75(63/9) | | \ | | \ | |
| GluA1i-TARP γ-2 | 93.89±<br>22.90(32/3) | | 127.22±<br>11.96(31/3) | | \ | | \ | |

Mean ( ± S.D.) values for constant of recovery from desensitization ( $\tau_{w, rec}$ ) from the recovery ratio curves, which were calculated using nonlinear one-phase exponential decay fitting. Currents were recorded in outside-out patch recordings (-60 mV, 10 mM glutamate; 100 ms) when performing two-pulse stimulation from cells transfected with GluA, GluA + ABHD6, GluA + TARP γ-2, and GluA + TARP γ-2 + ABHD6. The number of patches is given in parentheses.
<sup>a</sup>Significantly different from GluA.
<sup>b</sup>Significantly different from GluA + TARP γ-2.

**Table EV4.2.** Summary of P values for comparison of τ _w, rec_.

| Subunit | GluA+ABHD6<br>vs. GluA | GluA+TARP γ-2<br>vs. GluA | GluA+TARP γ-2+ABHD6<br>vs. GluA+TARP γ-2 |
| --- | --- | --- | --- |
| GluA1i | P>0.9999(-) | P<0.0001(***) | P=0.0092(**) |
| GluA1o | P>0.9999(-) | P<0.0001(***) | P=0.4938(-) |
| GluA2(Q)i-R | P>0.9999(-) | P=0.0325(*) | P>0.9999(-) |
| GluA2(Q)o-R | P>0.9999(-) | P=0.1988(-) | P>0.9999(-) |
| GluA2(Q)i-G | P>0.9999(-) | P=0.0014(**) | P=0.8413(-) |
| GluA2(Q)o-G | P>0.9999(-) | P>0.9999(-) | P>0.9999(-) |
| GluA2(Q)i-R-TARP γ-2 | P=0.075(-) | \ | \ |
| GluA1i-TARP γ-2 | P<0.001(***) | \ | \ |

P values for comparison of $\tau_{w, rec}$ of GluAs when co-transfected with/without γ-2 or/and ABHD6. The data were analyzed using Prism 8 (GraphPad Software), and each P value was adjusted to account for multiple comparisons.

**Table EV5.1.** Summary of GluA1i/GluA2(R)i-G receptors when co-transfected with/without γ-2 or/and ABHD6.

|  | **GluA1i/GluA2(R)i-G** |  | **GluA1i/GluA2(R)i-G + ABHD6** |  | **GluA1i/GluA2(R)i-G + TARP γ-2** |  | **GluA1i/GluA2(R)i-G + TARP γ-2 + ABHD6** |  |
| --- | --- | --- | --- | --- | --- | --- | --- | --- |
| Peak amplitude | 156.96 | ± | 65.75 | ± | 492.72 | ± | 67.45 | ± |
|  | 140.07(31/5) |  | 66.79(33/5)<sup>a</sup> |  | 454.74(34/5)<sup>a</sup> |  | 141.71(32/5)<sup>b</sup> |  |
| $\tau_{w, deact}$ | 1.04 | ± | 1.00 | ± | 2.20 | ± | 1.59 | ± |
|  | 0.46(31/5) |  | 0.30(33/5) |  | 0.98(34/5)<sup>a</sup> |  | 0.76(32/5)<sup>b</sup> |  |
| $\tau_{w, des}$ | 3.85 | ± | 3.24 | ± | 7.77 | ± | 4.27 | ± |
|  | 1.10(31/5) |  | 0.76(33/5) |  | 2.27(34/5)<sup>a</sup> |  | 1.53(32/5)<sup>b</sup> |  |
| $\tau_{rec}$ | 129.72 | ± | 263.71 | ± | 83.11 | ± | 164.42 | ± |
|  | 39.79(23/5) |  | 180.45(19/5)<sup>a</sup> |  | 28.70(23/5)<sup>a</sup> |  | 86.02(17/5)<sup>b</sup> |  |

Mean ( ± S.D.) values for peak amplitude, $\tau_{w, des}$ from the desensitization and $\tau_{w, deact}$ from the deactivation curves, and $\tau_{rec}$ from the recovery ratio curves. Currents were recorded in outside-out patch recordings (-60 mV, 10 mM glutamate; 100 ms / 1 ms / 100 two-pulse stimulation) without spermine in internal solution from cells transfected with GluA1i/GluA2(R)i-G, GluA1i/GluA2(R)i-G + ABHD6, GluA1i/GluA2(R)i-G + TARP γ-2, and GluA1i/GluA2(R)i-G + TARP γ-2 + ABHD6. The number of patches is given in parentheses.
<sup>a</sup>Significantly different from GluA1i/GluA2(R)i-G.
<sup>b</sup>Significantly different from GluA1i/GluA2(R)i-G + TARP γ-2.

**Table EV5.2.** Summary of P values for comparison of GluA1i/GluA2(R)i-G receptors when co-transfected with/without γ-2 or/and ABHD6.

|  | **GluA1i/GluA2(R)i-G**<br>**+ABHD6**<br>**vs.**<br>**GluA1i/GluA2(R)i-G** | **GluA1i/GluA2(R)i-G**<br>**+TARP γ-2 vs.**<br>**GluA1i/GluA2(R)i-G** | **GluA1i/GluA2(R)i-G**<br>**+TARP**<br>**γ-2+ABHD6**<br>**vs.**<br>**GluA1i/GluA2(R)i-G**<br>**+TARP**<br>**γ-2** |
| --- | --- | --- | --- |
| Peak amplitude | P=0.01(*) | P=0.007(**) | P<0.001(***) |
| $\tau_{w, deact}$ | P>0.99(-) | P<0.001(***) | P=0.03(*) |
| $\tau_{w, des}$ | P=0.43(-) | P<0.001(***) | P<0.001(***) |
| $\tau_{rec}$ | P=0.0109(*) | P=0.0076(**) | P=0.0011(**) |

P values for comparison of Peak amplitude, $\tau_{w, deact}$ , $\tau_{w, des}$ , $\tau_{rec}$ of GluA1i/GluA2(R)i-G when co-transfected with/without γ-2 or/and ABHD6. The data were analyzed using Prism 8 (GraphPad Software), and each P value was adjusted to account for multiple comparisons.

**Table EV6.1.** Summary of GluA2(R)i-G/GluA3(R)i receptors when co-transfected with/without γ-2 or/and ABHD6.

|  | **GluA2(R)i-G/GluA3(R)i** |  | **GluA2(R)i-G/GluA3(R)i + ABHD6** |  | **GluA2(R)i-G/GluA3(R)i + TARP γ-2** |  | **GluA2(R)i-G/GluA3(R)i + TARP γ-2 + ABHD6** |  |
| --- | --- | --- | --- | --- | --- | --- | --- | --- |
| Peak amplitude | 132.3<br>(19/3) | $\pm 55.74$ | 42.11<br>(18/3)<sup>a</sup> | $\pm 34.19$ | 514.2<br>(20/3)<sup>a</sup> | $\pm 171.1$ | 66.42<br>(18/3)<sup>b</sup> | $\pm 67.24$ |
| $\tau_{w, deact}$ | 1.128<br>(19/3) | $\pm 0.2640$ | 1.159<br>(18/3) | $\pm 0.3888$ | 2.325<br>(20/3)<sup>a</sup> | $\pm 0.8924$ | 1.284<br>(18/3)<sup>b</sup> | $\pm 0.3922$ |
| $\tau_{w, des}$ | 5.346<br>(19/3) | $\pm 1.203$ | 4.949<br>(18/3) | $\pm 2.735$ | 7.645<br>(20/3)<sup>a</sup> | $\pm 1.579$ | 5.160<br>(18/3)<sup>b</sup> | $\pm 3.823$ |
| $\tau_{rec}$ | 75.46<br>(18/3) | $\pm 89.31$ | 57.58<br>(18/3)<sup>a</sup> | $\pm 23.78$ | 81.31<br>(19/3)<sup>a</sup> | $\pm 35.58$ | 89.81<br>(18/3)<sup>b</sup> | $\pm 49.89$ |

Mean ( ± S.D.) values for peak amplitude, $\tau_{w, des}$ from the desensitization and $\tau_{w, deact}$ from the deactivation curves, and $\tau_{rec}$ from the recovery ratio curves. Currents were recorded in outside-out patch recordings (-60 mV, 10 mM glutamate; 100 ms / 1 ms / 100 two-pulse stimulation) without spermine in internal solution, from cells transfected with GluA2(R)i-G/GluA3(R)i, GluA2(R)i-G/GluA3(R)i + ABHD6, GluA2(R)i-G/GluA3(R)i + TARP γ-2, and GluA2(R)i-G/GluA3(R)i + TARP γ-2 + ABHD6. The number of patches is given in parentheses.
<sup>a</sup>Significantly different from GluA2(R)i-G/GluA3(R)i.
<sup>b</sup>Significantly different from GluA2(R)i-G/GluA3(R)i + TARP γ-2.

**Table EV6.2.**
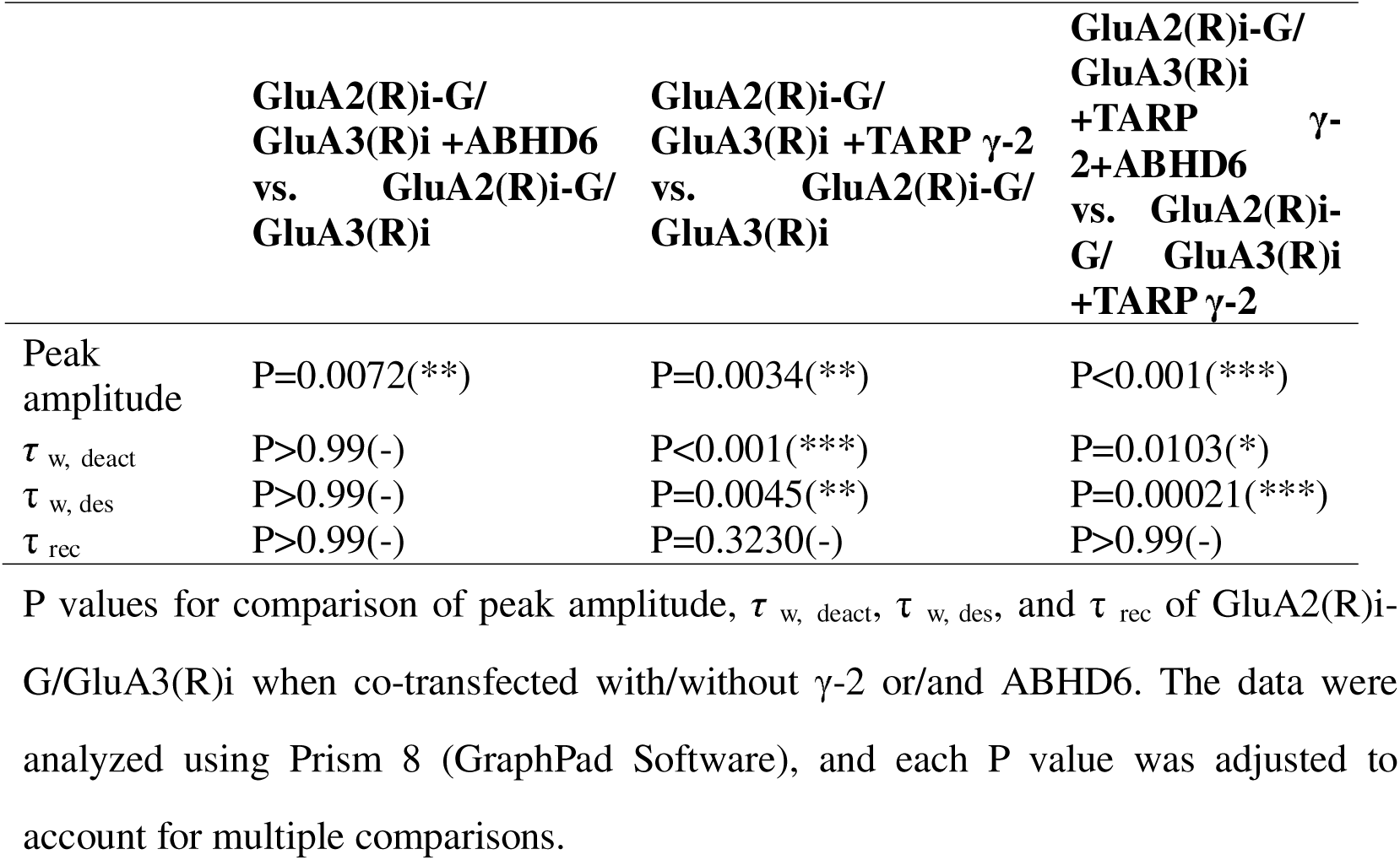
Summary of P values for comparison of GluA2(R)i-G/ GluA3(R)i receptors when co-transfected with/without γ-2 or/and ABHD6.

|  | **GluA2(R)i-G/**<br>**GluA3(R)i +ABHD6**<br>**vs. GluA2(R)i-G/**<br>**GluA3(R)i** | **GluA2(R)i-G/**<br>**GluA3(R)i +TARP γ-2**<br>**vs. GluA2(R)i-G/**<br>**GluA3(R)i** | **GluA2(R)i-G/**<br>**GluA3(R)i**<br>**+TARP γ-2+ABHD6**<br>**vs. GluA2(R)i-G/**<br>**GluA3(R)i**<br>**+TARP γ-2** |
| --- | --- | --- | --- |
| Peak<br>amplitude | P=0.0072(**) | P=0.0034(**) | P<0.001(***) |
| $\tau_{w, deact}$ | P>0.99(-) | P<0.001(***) | P=0.0103(*) |
| $\tau_{w, des}$ | P>0.99(-) | P=0.0045(**) | P=0.00021(***) |
| $\tau_{rec}$ | P>0.99(-) | P=0.3230(-) | P>0.99(-) |

P values for comparison of peak amplitude, $\tau_{w, deact}$ , $\tau_{w, des}$ , and $\tau_{rec}$ of GluA2(R)i-G/GluA3(R)i when co-transfected with/without γ-2 or/and ABHD6. The data were analyzed using Prism 8 (GraphPad Software), and each P value was adjusted to account for multiple comparisons.

**Table EV 7.1.** Summary of wild-type primary hippocampal neurons and ABHD6 knockout primary hippocampal neurons.

|  | WT | KO |
| --- | --- | --- |
| Peak amplitude | 243.2 ±264.5 (43/3) | 569.9±650.5 (44/3) |
| $\tau_{w, deact}$ | 2.163 ±0.8521 (43/3) | 2.616 ±1.044 (44/3) |
| $\tau_{w, des}$ | 4.406 ±2.284 (43/3) | 6.514 ±4.646 (44/3) |
| $\tau_{rec}$ | 133.3 ±122.0 (38/3) | 89.87 ±54.54 (43/3) |

Mean ( ± S.D.) values for peak amplitude, $\tau_{w, des}$ from the desensitization and $\tau_{w, deact}$ from the deactivation curves, and $\tau_{rec}$ from the recovery ratio curves. Currents were recorded in outside-out patch recordings (- 70 mV, 10 mM glutamate; 100 ms / 1 ms / 100 two-pulse stimulation) without spermine in internal solution from cultured WT or ABHD6 KO primary hippocampal neurons. The number of patches is given in parentheses.

**Table EV 7.2.**
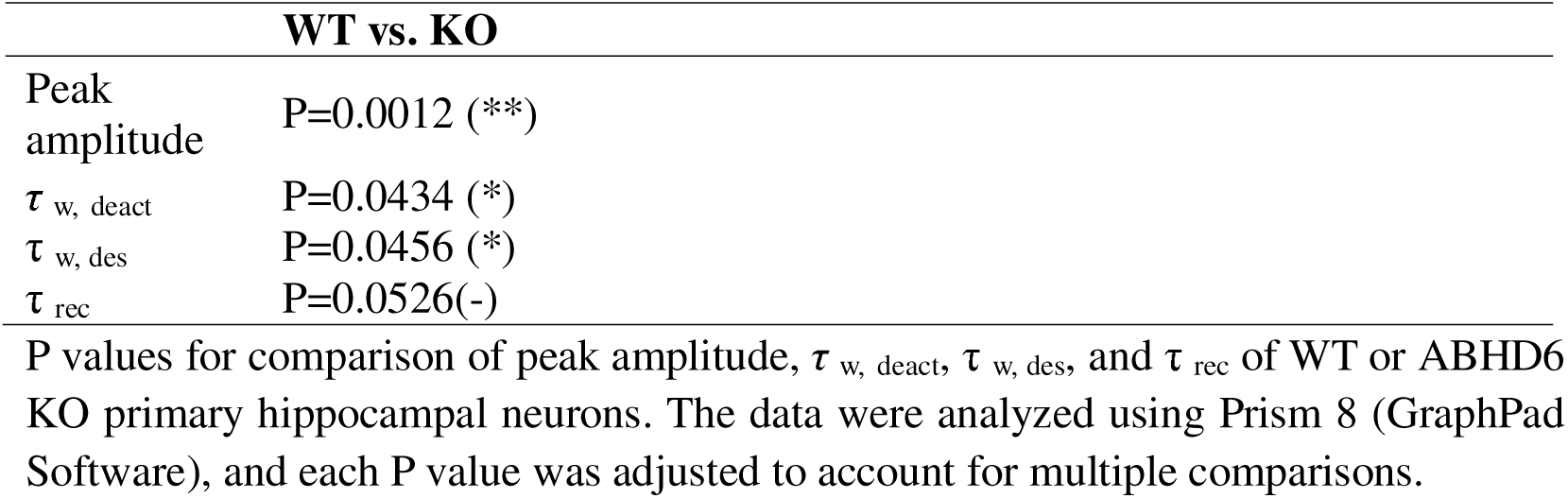
Summary of P values for comparison of wild-type primary hippocampal neuron and ABHD6 knockout primary hippocampal neuron.

**Table EV8.1.**
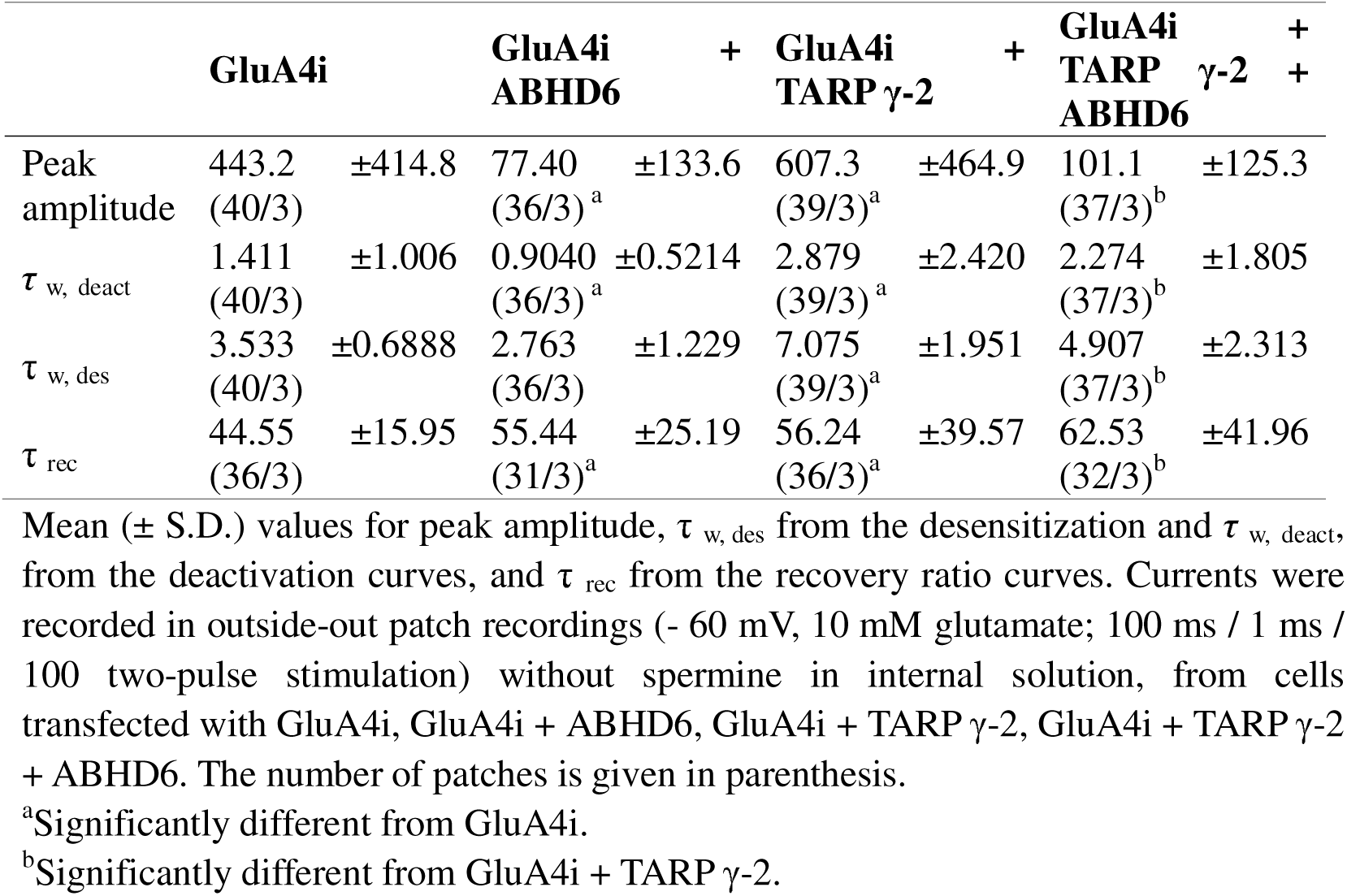
Summary of GluA4i receptors when co-transfected with/without γ-2 or/and ABHD6.

**Table EV8.2.** Summary of P values for comparison of GluA4i receptors when co-transfected with/without γ-2 or/and ABHD6.

| | GluA4i + ABHD6<br>vs. GluA4i | GluA4i + TARP γ-2<br>vs. GluA4i | GluA4i + TARP γ-2 + ABHD6<br>vs. GluA4i + TARP γ-2 |
| --- | --- | --- | --- |
| Peak amplitude | P<0.001(***) | P=0.5472(-) | P>0.9999(-) |
| $\tau_{w, deact}$ | P=0.2095(-) | P<0.001(***) | P=0.6753(-) |
| $\tau_{w, des}$ | P=0.1350(-) | P<0.001(***) | P=0.0006 (***) |
| $\tau_{rec}$ | P=0.1599(-) | P=>0.9999(-) | P=>0.9999(-) |

P values for comparison of Peak amplitude, $\tau_{w, deact}$ , $\tau_{w, des}$ , $\tau_{rec}$ of GluA4i when co-transfected with/without γ-2 or/and ABHD6. The data were analyzed using Prism 8 (GraphPad Software), and each p values was adjusted to account for multiple comparisons.

**Table EV9.1.**
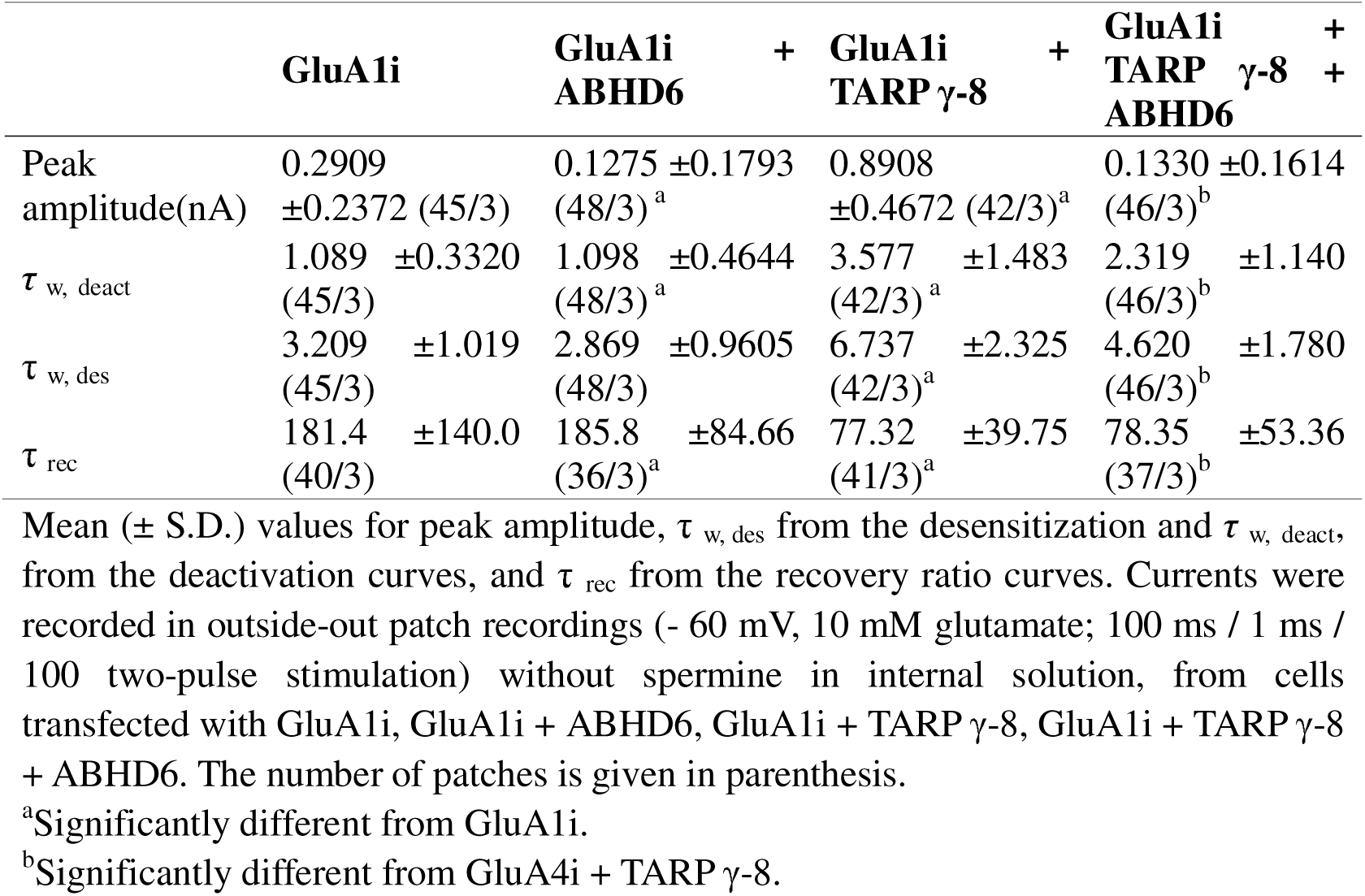
Summary of GluA1i receptors when co-transfected with/without γ-8 or/and ABHD6.

**Table EV9.2.** Summary of P values for comparison of GluA1i receptors when co-transfected with/without γ-8 or/and ABHD6.

|  | **GluA4i +ABHD6**<br>**vs. GluA4i** | **GluA4i +TARP γ-2**<br>**vs. GluA4i** | **GluA4i +TARP**<br>**γ-2+ABHD6**<br>**vs. GluA4i**<br>**+TARP γ-2** |
| --- | --- | --- | --- |
| Peak<br>amplitude | P=0.001(**) | P<0.001(***) | P<0.001(***) |
| $\tau_{w, deact}$ | P>0.9999(-) | P<0.001(***) | P=0.02(*) |
| $\tau_{w, des}$ | P>0.9999(-) | P<0.001(***) | P=0.001(**) |
| $\tau_{rec}$ | P>0.9999(-) | P<0.001(***) | P=>0.9999(-) |

P values for comparison of Peak amplitude, $\tau_{w, deact}$ , $\tau_{w, des}$ , $\tau_{rec}$ of GluA1i when co-transfected with/without γ-8 or/and ABHD6. The data were analyzed using Prism 8 (GraphPad Software), and each p values was adjusted to account for multiple comparisons.

## Notes

### Competing Interest Statement

The authors have declared no competing interest.

### Summary of Updates

In this version, Figure EV1, Figure 5, Figure EV5, Figure 6, Figure 7, and Figure EV7 have been updated. The new version includes additional data on the kinetic effects of ABHD6 overexpression on GluA2(R)i/GluA3(R)i heteromeric receptors, as well as the effects of ABHD6 knockout on the kinetics mediated by neuronal AMPA receptors.

